# Multi-Temporal Remote Sensing of Inland Surface Waters: A Fusion of Sentinel-1&2 Data Applied to Small Seasonal Ponds in Semiarid Environments

**DOI:** 10.1101/2024.03.03.583180

**Authors:** Francesco Valerio, Sérgio Godinho, Gonçalo Ferraz, Ricardo Pita, João Gameiro, Bruno Silva, Ana Teresa Marques, João Paulo Silva

**Author notes:** Corresponding author (<u>)∼</u>.

## Abstract

Inland freshwater resources in semiarid environments play a key role in maintaining ecological systems and supporting human development. Space-based remote sensing spatiotemporal data have emerged as a new paradigm for understanding ecohydrological processes and trends, particularly in water-stressed areas. However, comprehensive cataloging is still lacking, especially in semi-arid regions and for small-sized water bodies (i.e., ponds), which are often overlooked despite their ecological relevance. In this study, high-resolution optical and radar Sentinel data (Sentinel-1 and Sentinel-2) were used to construct Sentinel-1&2-based local surface water (SLSW) models, to infer surface water occurrence and extent. To assess the reliability of this model, the results were compared with verification data, and separately with Landsat-based global water (LGSW) models. Three distinct semiarid regions were selected in SW Iberia, within a Mediterranean climate, each encompassing special protection areas for conservation and subjected to marked seasonality and bioclimatic changes. Surface water attributes were modeled using Random Forests for SLSW time series forecasting, which included the period from January 1, 2020, to December 31, 2021. During this period, the completeness of the archived information was compared between SLSW and LGSW, considering both intra-annual and inter-annual variations. The predictive performance of these models was then compared for specific periods (dry and wet), and each was independently validated with verification data. The results showed that SLWM achieved satisfactory predictive performances in detecting surface water occurrence (*μ*≈72%), with far greater completeness and reconstructed seasonality patterns compared to LGSW. The relatedness between SLSW and LGSW was stronger during wet periods (R^2^=0.38) than dry periods (R^2^=0.05), and SLSW related much better with the verification data (R^2^=0.66) than when compared to LGSW (R^2^=0.24). The proposed SLSW approach may therefore provide advantages in the delineation of dynamic surface water characteristics (occurrence and extent) in very small-sized water bodies (i.e., <0.5 ha), allowing for uninterrupted surface water time series forecasting at high spatiotemporal detail, and over extensive areas. Given the water constraints in semiarid regions and water resources vulnerability to climate change, our results show high potential for supporting a variety of activities underlying rural development and biodiversity conservation. Additionally, the socio-ecological applications of this research may help identify surface water anomalies (e.g., drought events) and enhance sustainable water supply governance, a particular priority in climate change hotspots.

**Highlights:**

- Surface water occurrence and extent was modeled across three semiarid regions
- Sentinel-1&2 data was compared with Landsat for characterizing very small water bodies
- Models based on Sentinel-1&2 resulted in a satisfactory classification precision
- Very high series completeness across seasons was found using Sentinel-1&2 data
- Sentinel-1&2 data was more reliable than Landsat when compared to verification data

## 1. Introduction

Water is essential for life on Earth and human development (Hoekstra, 2009). Given the contributes that inland freshwater systems play in supporting biological cycles and carbon storage, and the risks posed by rising human needs leading to water contamination and overuse, a balance between environmental sustainability and public water management is crucial (Millennium Ecosystem Assessment, 2005; Gleick, 1998). Sustainability concerns further arise because inland waters are a limited natural resource with scarce geographical representation (∼3%) on the Earth’ biosphere (Downing et al., 2006), of which only a portion (∼35%) represents a globally disposable renewable resource (Millennium Ecosystem Assessment, 2005). The mounting evidence of water scarcity in many regions and associated socio-economic and ecological impacts, has progressively pressed for advancements in freshwater detection and monitoring strategies. Such advancements have the potential to offer guidance on how to mitigate stressors and overexploitation within natural freshwater systems and to enhance resilience towards climate change (Millennium Ecosystem Assessment, 2005; Erwin, 2009). As a result, the spatiotemporal dynamics of global and local freshwater systems have emerged as a topic of prime interest and a key Sustainable Development Goal (SDG; UN, 2022).

Spatiotemporal information from Earth Observation Satellites (EOS), comprising optical and radar sensors, has opened a new era for assessing qualitative and quantitative dynamic attributes of freshwater resources (Bijeesh and Narasimhamurthy, 2020; Gholizadeh et al., 2016; Petropoulos et al., 2015). Data covering large geographic areas represents an important milestone for sustainable water management, especially in water-stressed areas such as arid and semiarid regions, which cover more than 30% of Earth’ land surface (Wickens, 1998). Due to the high intra- and inter-annual variability of these ecosystems, resource management programs require estimates of surface water and systematic site monitoring at an appreciable level of detail (Scholes 2020; Wickens 1998). Because water-related catalogues are often insufficient, understanding ecohydrological processes is traditionally achieved via remote sensing information, even though the selection of a sensor and type of application are typically context-specific (Bijeesh and Narasimhamurthy, 2020; Gxokwe et al., 2022). Most research has been devoted to inferring water characteristics and dynamics over the long term through quantitative remote sensing methods. Among these, EOS multi-spectral information from MODIS (or Moderate Resolution Imaging Spectroradiometer) stands out due to its exceptional temporal resolution, but limited spatial resolution by today’ standards (250m), therefore being more suitable for large water reservoirs (Klein et al., 2017). In this sense, it is worth noting the substantial progress made in employing advanced instruments, such as public sensors affiliated with Landsat missions, and their deriving products, that have fundamentally advanced surface water mapping (Pekel et al., 2016; Zou et al., 2018). These advancements ensure higher spatial, radiometric, and spectral resolution, making Landsat a preferential candidate for water detection, particularly for smaller water bodies that are more difficult to detect (Balázs et al., 2018; Halabisky et al., 2016; Tulbure et al., 2017).

Small-sized water bodies (and similar systems like riparian ecosystems) are consistently overlooked in multiple research areas, including remote sensing, despite their critical importance for numerous ecosystems (Céréghino et al., 2008; Olmo et al., 2022; Wang et al., 2022). Possible causes hindering scientific progress are attributable to the difficulties in conducting complete ground surveys of such water bodies, which often appear isolated and scattered, thus requiring high manpower for their identification, particularly those of limited dimensions (< 5 ha; De Meester et al., 2005). Yet, despite being largely under-censused and rarely considered in large-scale inventories and mapping programs, small-sized water bodies are hypothesized to be more frequent than those large (i.e., lakes), and to cover a far greater fraction of Earth’ surface (Downing et al., 2006; Olmo et al., 2022). Formerly known as ‘ponds’, these elements are considered to play a relevant role in human well-being, offering secondary products (e.g., food, timber, and fuel), but also supporting fishing and agricultural activities (crops and livestock), as well as a plethora of ecosystem services including nutrient cycling, soil formation, and pollination (Millennium Ecosystem Assessment, 2005; Céréghino et al., 2008). They are also distinguished by different ecological processes, with greater heterogeneity and amplitudes of environmental and biotic conditions than larger water bodies (Casas et al., 2012; Olmo et al., 2022; Williams et al., 2004), while classified frequently as biodiversity hotspots (Williams et al., 2004). Also, it is observed that different types of ponds, either temporary or permanent, tend to host varying levels of species richness and abundance (Céréghino et al., 2008; Sebastián-González & Green, 2014; Williams et al., 2004). Importantly, freshwater habitats provide refuge for about 10% of known species, notwithstanding they are classified as among the most endangered ecosystems worldwide (Reid et al., 2019). In semiarid areas, environmental conditions are expected to worsen due to widespread extreme heat events, desertification processes, increased salinity, and agricultural intensification, with implications for water deficit anomalies (Diffenbaugh et al., 2007; Reid et al., 2019; Wickens, 1998).

Despite the importance of small water bodies, they are routinely misrepresented in traditional satellite-derived products (e.g., land cover; Coops and Wulder, 2019; Valerio et al., 2020). This may also occur for specific water-related products such as those from Landsat, owing to limitations for comprehensive assessments of very small-sized bodies (e.g., <0.5 ha; Sethre et al., 2005; Cordeiro et al., 2021). Limitations include the identification of objects smaller than the sensor grain (30m) and seasonal waters during cloudy periods, requiring improvements for EOS-based approaches (Ozesmi et al., 2002; Pekel et al., 2016). From this perspective, the affordable high-quality data from Sentinel missions have gained ground as a step forward over Landsat-derived data towards small water bodies’ characteristics delineation and mapping, given a better compromise between spectral, temporal, and spatial resolution (Du et al., 2016; Wang et al., 2022). Mainstream methods for mapping surface water bodies traditionally relied on single bands or spectral indices (Bijeesh and Narasimhamurthy, 2020; Du et al., 2016; Zhou et al.,2017). Histogram thresholding methods performed on EOS imageries have also been used to generate water and non-water classification maps, yet these have intrinsic limitations when dealing with low and high reflectance values (e.g., Peña-Luque et al., 2021). Alternatively, more sophisticated approaches relied on unsupervised and supervised water classification, with the latter increasingly adopting machine learning algorithms such as Random Forests given the high model robustness (Bangira et al., 2019; Bijeesh and Narasimhamurthy, 2020; Jiang et al., 2022). Another emerging procedure receiving increasing attention emphasizes the importance of employing multi-temporal imagery, notably sourced from missions such as Landsat and Sentinel. These missions have been instrumental in elucidating seasonal variations in water dynamics (Ozesmi et al., 2002; Vanderhoof et al., 2023; Tulbure et al., 2016; Tulbure et al., 2022). Regarding the Copernicus satellite constellation, recent water-related research efforts have increasingly focused on Sentinel-2 for classification purposes (Jiang et al., 2022; Radoux et al., 2016), involving the use of Sentinel-1 imageries (Huang et al., 2018; Shen et al., 2019), as well as combining both Sentinel-1 and Sentinel-2 data (Bioresita et al., 2019; Chen & Zhao, 2020; Mahdianpari et al., 2018; Vanderhoof et al., 2023). The fusion of multiple sensors, notably from Sentinel-1&2, may be especially promising when dealing with complex and seasonal small water bodies, considering that local events (e.g., drought, turbidity, anoxia, aquatic plants) can alter water spectral signature (Peña-Luque et al., 2021). The combination of Sentinel-1&2 has the potential to enhance the monitoring of surface water distribution and seasonality, particularly in small water bodies, though this issue remains poorly investigated in current literature. It is also unclear whether Sentinel-derived products offer more promising results compared to other high-resolution approaches (e.g., Landsat) for detecting and quantifying water surface in semiarid environments.

In this study, a data-fusion approach using Sentinel-1&2 information in semiarid environments is proposed to address data gaps and improve predictions on the spatiotemporal trends of dynamic surface water, specifically targeting open and very small water bodies. To assess the effectiveness of the approach, water bodies were mapped, and the Sentinel-1&2-based local surface water (SLSW) predictions were compared with those from Landsat-based global surface water (LGSW) (Pekel et al., 2016). Specifically, we aim to: (*i*) identify the best Sentinel-1&2-derived metrics to detect surface water occurrence, and understand multi-spectral reflectance and backscattering responses involved; (*ii*) utilize SLSW predictions to characterize the intra- and inter-year (2020-2021) dynamic trends (seasonality patterns) in surface water occurrence and extent (percentage) in three semiarid regions; and (*iii*) compare the SLSW archives completeness with that derived from the LGSW (Pekel et al., 2016), and evaluate the agreement between the estimated water extent from both data sources (Sentinel and Landsat), with a specific focus on the wet season (January-April) and the dry season (June-September) using a verification dataset.

## 2. Materials and methods

### 2.1. Study areas

Three study areas in the Mediterranean biome in southwestern Iberia were chosen to test the reliability of Sentinel-1&2-based data for accurate water surface mapping (Figure 1a): two in Baixo Alentejo (southern Portugal; PT1, PT2), and one in Extremadura (south-eastern Spain; SP1) (Figure 1a; 1b), wherein small water bodies were detected and mapped (Figure 1c).

**Figure 1.**
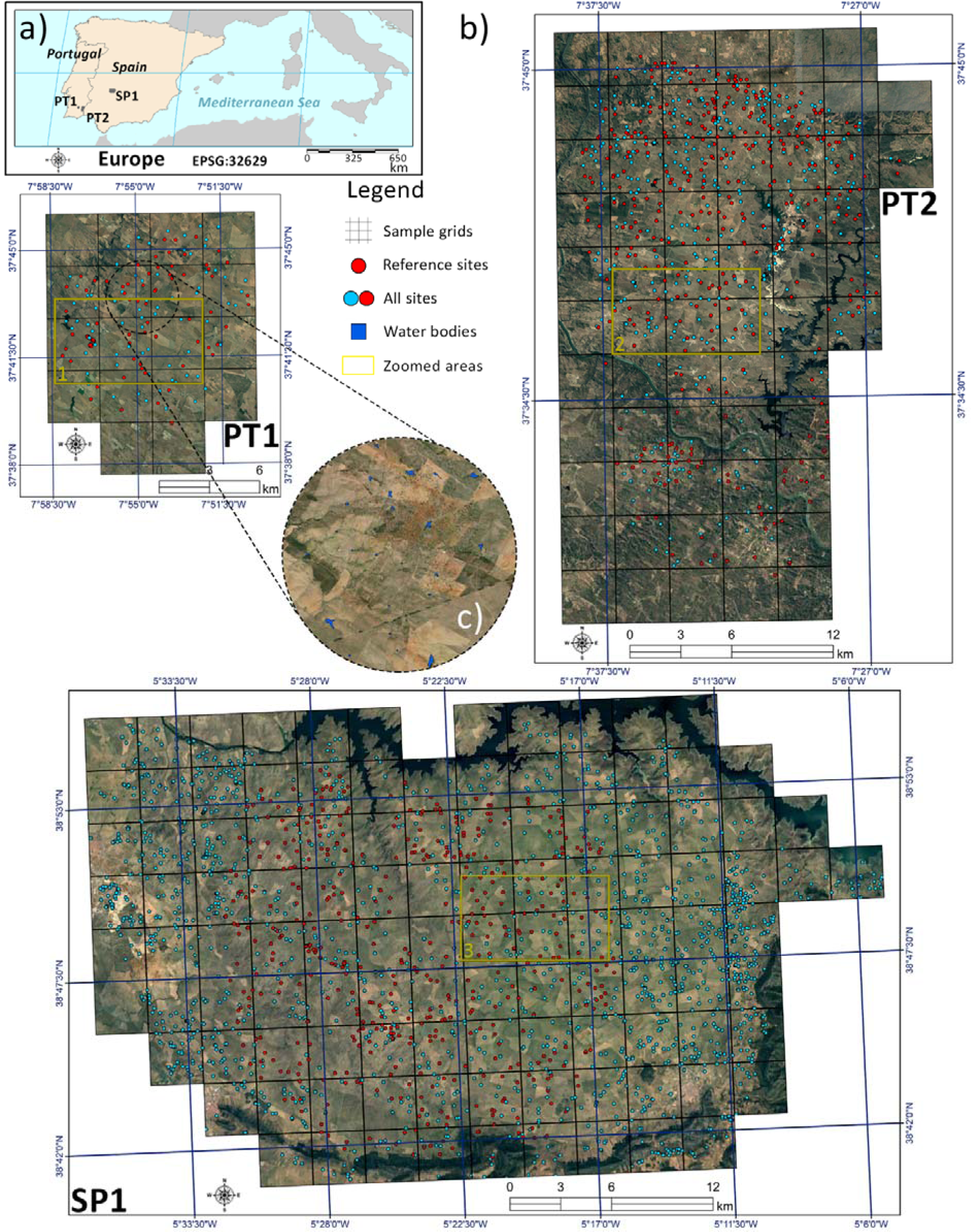
– The geographic locations of the study areas (a), and pertaining panels showing the distribution of water body sites as centroids (b), with red ones used as a reference for assessing data-fusion model performances. Panels of study areas are layered by an RGB color composite of Sentinel-2 overlapped by sample grids, each of 3×3km. Illustrative water bodies are represented in blue (c). The regions delimited in yellow (1, 2, and 3) serve the orienting purpose of identifying geographic locations depicted in Figure 8.

General similarities between the areas include climatic, topographic, and environmental conditions. In particular, the wet and dry seasons manifest a pronounced climatic variation, inducing severe influences on the water cycle. All areas are characterized by a semiarid climate with drought-hot summers and cold-wet winters, with most of the precipitation occurring from autumn to spring (AEMET and IMP, 2011). Typical weather parameters within the three areas were calculated for the period between 2012 and 2022 using the E-OBS daily gridded meteorological data (Cornes et al., 2018; Copernicus Climate Data Store, 2020), resulting in an annual average temperature of 17.42 °C (±SD = 0.52), a maximum temperature of 23.98 °C (±SD = 0.76), and an annual average precipitation amount of 450.69 mm (±SD = 122.24). Besides, soils in these areas are generally poor and, consequently, have limited water-retention capacity. As a result, water bodies are mostly very small (*Mdn*≈0.031 ha; Figure SM1), and sparsely distributed (Figure 1b). The three areas are characterized by plains and gentle slopes, dominated by pastures and extensive cereal agriculture (Gameiro et al., 2020). Each area is located within a Special Protection Area (Natura 2000 Network): Castro Verde for PT1, Vale do Guadiana for PT2, and La Serena y Sierras Periféricas for SP1. Their conservation value also resides in holding High Nature Valued Farmlands, with high levels of biodiversity and a community of key endangered species (e.g., *Tetrax tetrax*; *Otis tarda*) (Silva et al., 2024). Yet, those ponds are also included within a climate change hotspot, where drought events and water deficit anomalies are expected to increase (Samaniego et al., 2018).

### 2.2. Study design

The study design is comprehensively depicted in Figure 2, including a series of distinct steps, from water bodies’ inventories and remote sensing data collection and processing, to the subsequent modelling stages for estimating surface water occurrence and extent as high-frequency time series.

**Figure 2.**
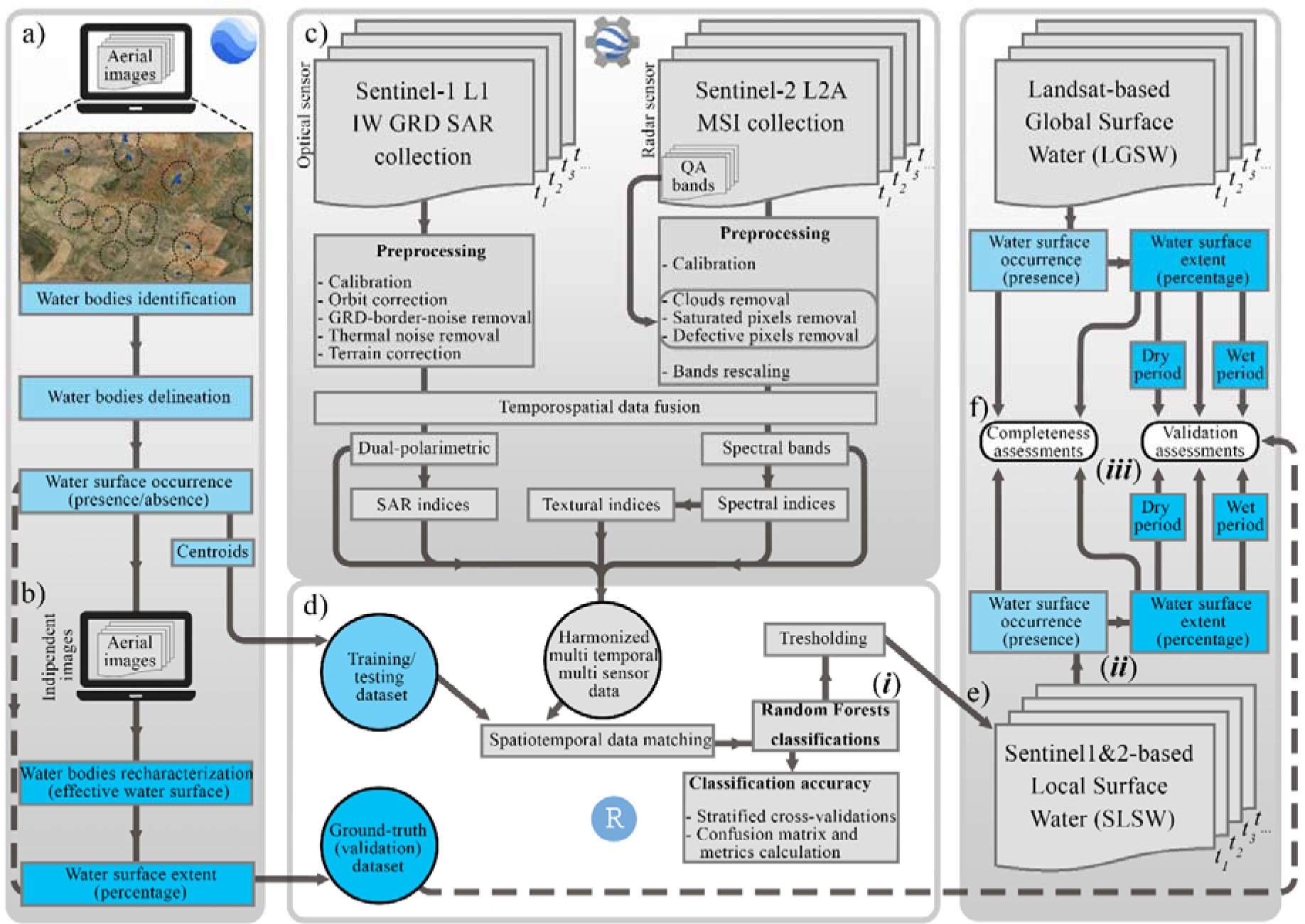
– Framework diagram depicting the analytical steps along four main panels, where arrows represent the operational progressions. The left panel illustrates how water bodies’ datasets were prepared, namely surface water occurrence (a) and extent (b), whilst the central-upper panel (c) shows how satellite (Sentinel-1&2) image time series were pre-processed and how variables were calculated. The central-lower panel (d) indicates how data was compiled for subsequent analyses to respond to the first objective (*i*). The right panel depicts the second (*ii*) and third (*iii*) objectives following the characterization of the SLSW time series (e) and the comparative assessment with the LGSW time series (f).

### 2.3. Data acquisition and pre-processing

#### 2.3.1 Water bodies data collection

The survey and characterization of water bodies were conducted within a GIS environment through photointerpretation of high-resolution imageries (Google Earth software; Google) (Figure 2a), during both dry (August and September) and wet period conditions (November and March), from 2017 to 2021. The decision to extend the survey period beyond the specific timeframe considered to characterize surface water occurrence and extent (2020-2021) was necessary to comprehensively cover all study areas under consideration (Table SM1). The margins (shorelines) of water bodies were vectorized (Figure 1c) based on the maximum water capacity, determined by observing the separation of evident flood-prone shores from the surrounding landscape. This criterion included the transitional zone (littoral zone), but excluded the vegetative buffer zone (Casas et al., 2012). Eligible water bodies included dammed ponds, reservoirs, natural and semi-natural ponds (temporary, permanent, semipermanent, and farm ponds), while excavations, pools, and water treatment systems were ignored (Casas et al., 2012; Céréghino et al., 2008). A total of 3366 water bodies were vectorized (centroids in Figure 1b), from which 932 were used as a reference dataset (surface water occurrence) for further analysis. This reference dataset was selected using proportional stratified random sampling by parameterizing polygon frequencies within blocks of 3 × 3 km grids (Figure 1b) as population ‘*strata’* (Cochran, 1977). Water presence/absence characteristics (water and non-water classes) were included as an attribute to these polygons, jointly with an ID, and a timestamp (YYYY/MM/DD) reflecting the acquisition date of the imagery used.

Additionally, a separate ‘verification’ dataset was created to validate the surface water extent model predictions (Figure 2b). This dataset aimed to represent the effective surface water occupied within each water body and was created by randomly selecting 606 polygons from the original dataset (∼18%), which underwent a recharacterization of water body margins following imageries interpretation for an independent year (May 2021; Table SM2). The area comparison between the original dataset (maximum capacity water surface) and the verification dataset (effective surface water) ensured the quantification of the relative surface water extent (surface water percentage in water bodies), which was included as an attribute field for each water body.

#### 2.3.2 Sentinel-1&2 data pre-processing

The remote sensing data were derived from the Sentinel-1&2 constellations designed by the European Space Agency (ESA). The Sentinel-1 satellite carries a phase-preserving dual-polarization synthetic aperture radar (SAR) instrument (Torres et al., 2012), while the Sentinel-2 satellite carries a multispectral instrument (MSI) (Drusch et al., 2012). Satellites have a swath width of 250 km for Sentinel-1, and 290 km for Sentinel-2. For the analysis, the Level-2A product from the Sentinel-2A and Sentinel-2B generations was used, while, for Sentinel-1A and Sentinel-1B, the Level-1 Ground Range Detected (GRD) product was selected, with the Interferometric Wide (IW) swath as a baseline acquisition modality for both ascending and descending orbits. The Sentinel-1&2 imageries ingestion and pre-processing analyses were conducted using Google Earth Engine (GEE) (Figure 2c), a cloud computing platform able to communicate with different repositories and rapidly process large amounts of geospatial information (Gorelick et al.,2017). For the Sentinel-2 data, the following preprocessing steps (Figure 2c) were performed: the MSI data was converted to bottom-of-atmosphere reflectance, orthorectified, and subjected to atmospheric corrections. Surface reflectance values, expressed on a scale of 0 to 1 and holding physical significance, were derived by applying a rescaling factor to Level-2A digital numbers (DNs). Cloudy objects were removed using the quality assessment (QA) QA60 band, following Mahdianpari et al., (2018). Bitwise operations were applied along the time series to mask out opaque and cirrus clouds, as well as cloud shadow and noisy saturated pixels. However, dark area pixels were not intervened to avoid losing the signature of water bodies with appreciable dark bottoms (Peña-Luque et al., 2021), which can be caused by anoxic sediments commonly found in poorly mobilized waters (McKenzie et al., 2001). For Sentinel-1 data, the processing steps (Figure 2c) included: orbit correction, GRD border and thermal noise removal, radiometric calibration, and terrain correction (orthorectification) converting the sigma nought (σ°) backscatter coefficient into decibels (dB).

Another step for the two collections involved a spatiotemporal data optimization due to the heterogeneity of the imageries resulting from different satellite origins, orbits, acquisition dates, grain sizes, and frequency in sites revisitation (3 to 6 days) (Drusch et al., 2012; Peña-Luque et al., 2021; Radočaj et al., 2020). A temporospatial data fusion (*sensu* Liu et al., 2018) (Figure 2c) was applied to align the two collections (co-registration; e.g., Chen & Zhao, 2022), followed by a routinely compositing of the imageries into 15-day time intervals at high resolution (10m). High-quality imageries that previously underwent masking operations from QA bands were here composited at each time interval to fill missing information. An 8-meter resampling procedure was further employed for pixels arrangement purposes, with the specific aim of improving pixel alignment along non-uniform water polygon edges, while reducing pixel surface areas extending beyond water body areas. Therefore, only polygons larger than 0.0064 ha, corresponding to a pixel of 8 × 8 m, were considered for the analysis. These sub-steps resulted in a harmonized dataset comprising 48 high-resolution, high-quality imageries spanning from 2020 to 2021.

### 2.4 Environmental factors

To assess the ability of Sentinel-1&2-derived metrics to flag the occurrence of surface water (i.e., probability of occurrence) (objective *i*), bands from Sentinel-1&2 were included (Figure 2c). From the Sentinel-2 reflectance bands, those with a native spatial resolution of 60m (cirrus band, coastal aerosol, and water vapor) were discarded, resulting in 10 high-resolution bands. These bands covered the visible spectrum (blue, green, red), near-infrared spectrum (NIR1&2, Red-edge1&2&3), and short-wave-infrared spectrum (SWIR1&2) (Drusch et al., 2012). The Sentinel-1 data was represented by the C-band SAR with dual-polarized channels: a co-polarized vertical transmit and vertical receive (VV), and a cross-polarized vertical transmit and horizontal receive (VH) (Attema et al., 2008).

Following the resampling procedure previously described, spectral and SAR indices were generated separately from Sentinel-1&2 data (Figure 2c), and integrated within the 2-year time series. For Sentinel-2, water-related indices included the normalized difference moisture index (NDMI; Wilson and Sader, 2002) and the normalized difference water index 2 (NDWI2; Gao, 1996; Serrano et al., 2000), each alternatively incorporating the NIR1 or NIR2 band (NDMI_N1, NDMI_N2, NDWI2_N1, NDWI2_N2). Additionally, the Sentinel-2 water index (SWI; Jiang et al., 2020) was computed (Table 1). Several vegetation indices were also considered such as the normalized difference vegetation index (NDVI; Rouse et al., 1974), and the modified soil-adjusted vegetation index 2 (MSAVI2; Richardson and Wiegand, 1977), both calculated with either the NIR1 or NIR2 band (NDVI_N1, NDVI_N2, MSAVI2_N1, MSAVI2_N2). The normalized difference index 4 and 5 (NDI45; Delegido et al., 2011) was also calculated (Table 1). Textural indices were also computed (Figure 2c), specifically to address complex surface conditions often found in small water bodies (Bangira et al., 2019). This involved the calculation of the multi-direction (0°, 45°, 90°, and 135°) grey level co-occurrence matrix (GLCM) using second-order statistics (Haralick et al., 1973). The pattern signature complexity was measured through the combinations of pixel brightness values (grey levels) within a parameterized kernel. A kernel size of 3 × 3 pixels was applied, and the textural indices computed from the NDVI images were contrast, variation, homogeneity, and mean (GLCM-C, GLCM-V, GLCM-H, GLCM-M) (Table 1). For Sentinel-1, water-related SAR indices included the VH and VV backscatters cross ratio (VHVVR or VH/VV; Alvarez-Mozos and Gonzalez-Audicana, 2021), the normalized difference polarization index (NDPI; Cao and Zheng, 2008), the dual-polarized radar vegetation index VV (RVI; Nasirzadehdizaji et al., 2019), as well as the associated indices normalized VH index (NVHI) and normalized VV index (NVVI) (McNairn and Brisco, 2004). These indices were computed to improve surface water detection and provide information on landscape biophysical attributes such as soil and vegetation moisture (Table 1) (Bangira et al., 2019; Huang et al., 2018; Mayer et al., 2021).

**Table 1.**
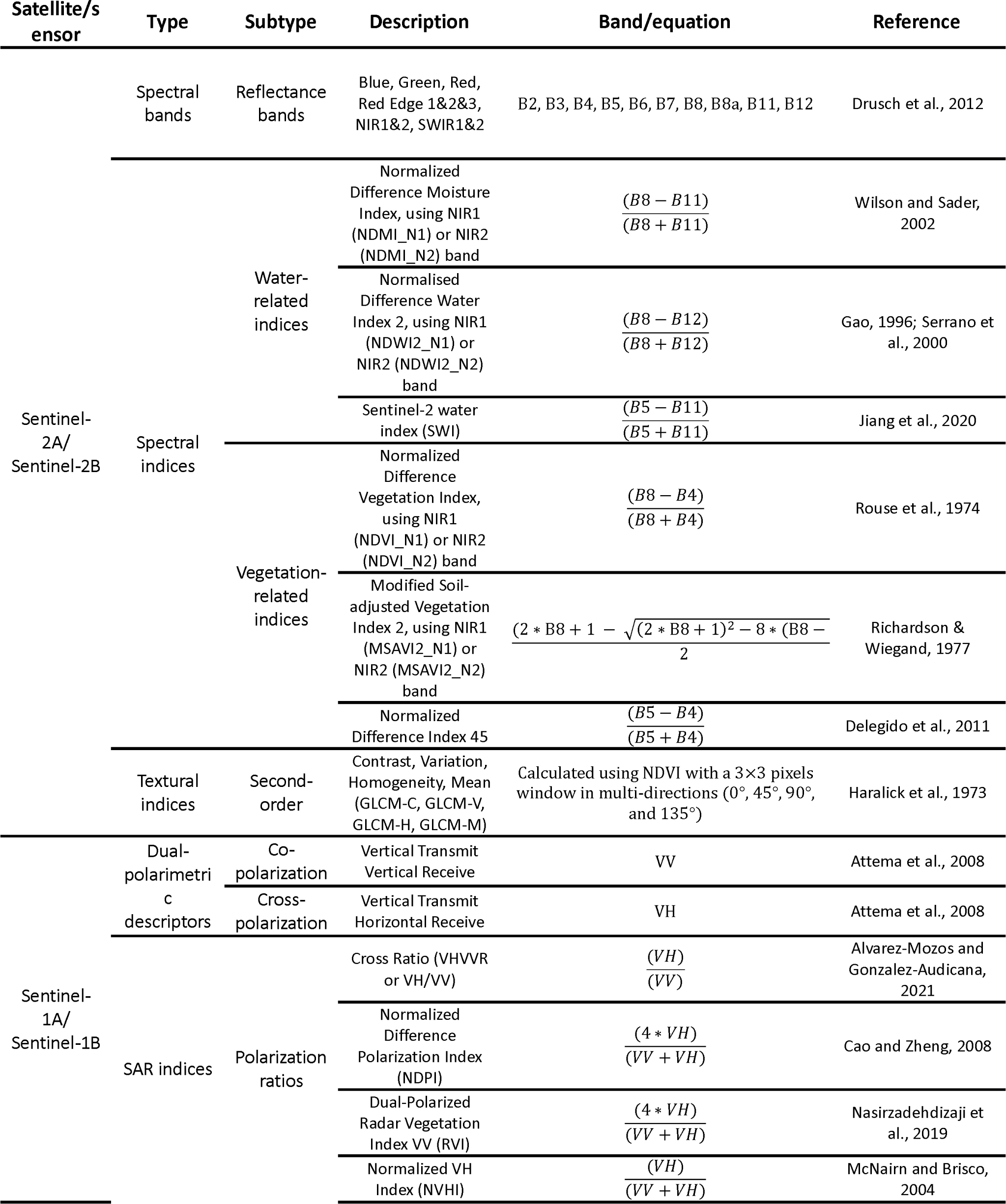
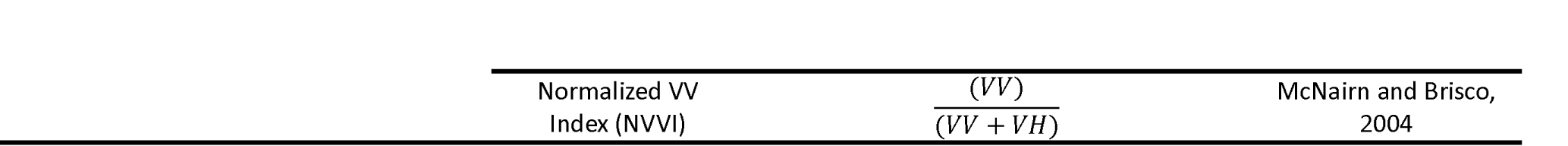
*–* The portfolio of spectral and SAR bands with derived indices selected for detecting surface water in small water bodies.

### 2.5 Surface water occurrence

Prior analysis, the vector dataset was converted into centroids (Figure 2a), assuming that central pixels were the best indicators of water presence. The ID and timestamp fields from the centroids served as input data for various sequential operations in the web-based tool *GEE_xtract* (Valerio et al., 2024), ensuring the extraction of environmental data from centroids with a high spatial match, while providing reasonable temporal alignment in the time series (15-days frequency) forecasting. To determine water presence (occurrence) within polygons, a binary classification was selected to discriminate between water and non-water classes (response variable), with calculated environmental factors used as explanatory variables. The Random Forests (RF; Breiman, 2001) algorithm was used as a supervised machine learning method (Figure 2d), given its robustness in extracting information about water bodies from space-based remote sensing data (Bangira et al., 2019; Balázs et al., 2018; Jiang et al., 2022; Peña-Luque et al., 2021). RF is an ensemble learning method of bagging and bootstrapping, where a forest of decision trees is built and trained by splitting at each independent tree node both response and explanatory variables. As each decision tree is trained on a subset of the original datapoints, the remaining datapoints, a.k.a. out-of-bag (OOB) samples, can be used to obtain the OOB error for that particular tree, a metric commonly used for model validation (Breiman, 2001). Furthermore, while RF is capable of dealing with high-dimensional information (Jiang et al., 2022), environmental variables with a Pearson’ correlation coefficient |r|< 0.9 (Millard and Richardson, 2015) were retained. To increase the comparability of our results, the variable selection among highly correlated variables was expert-based according to main recent advances in water surface mapping using Sentinel-derived data (Chen & Zhao, 2022; Du et al., 2012; Mayer et al., 2021; Peña-Luque et al., 2021; Radoux et al., 2016). Hence, our initial dataset resulted in 14 eligible variables for further analysis (Figure SM2). The RF algorithm was then configured with a node-splitting parameter equal to the square root of the total of selected variables, and a total of 2000 trees (Valerio et al., 2020). For the analyses, it was considered the reference dataset of 932 water body observations (∼28% of the total dataset), sub-divided into water (503 observations) and non-water classes (429 observations). The multivariate RF models were assessed through cross-validations (Figure 2d). The data, divided into 70% for training and 30% for testing, was stratified into 5 k-folds based on spatial blocks of 3 × 3 km (Figure SM3). This means that each of the five RF models was trained on a distinct geographic spatial block, though their extrapolations were applied to the entire areas covered by water bodies and then averaged. This approach aimed to minimize autocorrelation problems and maximize model extrapolation power (Valavi et al., 2018). The RF model’ performance (classification accuracy; Figure 2d) was primarily assessed using a confusion matrix (Stehman, 1997), computed for each of the stratified cross-validations. From these matrices, several accuracy metrics were averaged: Area Under the Curve (AUC; Swets, 1988), Sensitivity (proportion of occurrences correctly classified), Specificity (proportion of absences correctly classified; Gašparović and Singh, 2022), Accuracy (Bangira et al., 2019), F1 score (Mayer et al., 2021), and Matthews’ Correlation Coefficient (MCC; Baldi et al., 2020). The importance of variables for classification (water and non-water classes) and for surface water occurrence detection was determined using the Gini index (or Mean Decrease Impurity) (Breiman, 2001), which measures the effect of each variable in explaining surface water occurrence. To select significantly influential variables, an importance cutoff was implemented and derived by the averaged importance scores including all variables (Valerio et al., 2020). Then, the response curves were examined through partial dependence plots.

The RF multivariate analyses were repeated along the previously selected time series (48-time intervals X 5-fold cross-validation) to infer surface water occurrence probability, at each time interval with retained variables (14), for a total of 672 variables (14 × 48). The continuous classification probability was then subjected to a thresholding procedure to predict ‘water’ and ‘non-water’ pixels within the original polygons representing maximum water capacity. The optimal threshold (P=0.51) was determined using five combined techniques (further details in SM1.1). The surface water occurrence was computed using polygons containing at least one pixel flagged as with water, and then compared proportionally to those lacking any water pixels.

### 2.6 Surface water extent

To measure the surface water extent, the cumulative area from the predicted surface water occurrence was calculated within each polygon, and compared to the original polygon area, at each time interval. To account for overlaps of surface water pixels that extend beyond the margins of the polygons, and to deal with occasional flooding events (Tulbure et al., 2022), an 8-m buffer distance reflecting the raster grain was implemented around each polygon. All these steps were repeated along the modeled time series, referred to as SLSW (Sentinel-1&2-based Local Surface Water), enabling the investigation of the temporal trends in surface water occurrence and extent (objective *ii*; Figure 2e).

### 2.7 Reliability assessment of the surface water model

The reliability of our modeling approach was tested through a high-quality external model and with the independent verification dataset. The JRC Monthly Water History (v1.4) by Pekel et al. (2016), a Landsat-based high-quality model of surface water occurrence, was used as an external validation model. This product consists of 454 modeled imageries (from 4,716,475 scenes) spanning from 1984 to 2022. It provides globally available pixel-level surface water information derived entirely from Landsat 5, 7, and 8 sensors at high resolution (Pekel et al., 2016). A monthly time series coincident with the SLSW period was retained and referred to as LGSW (Landsat-based Global Surface Water; Figure 2f). The LGSW underwent the same steps as SLSW to enable the comparison between models in terms of archive completeness (objective *iii*; Figure 2g). The SLSW time series forecasting in water surface occurrence and extent were further scrutinized following the comparison with trends derived by the E-OBS gridded data (Cornes et al., 2018), with a focus on parameters such as mean and maximum temperature, precipitation amount, relative humidity, and surface shortwave downwelling radiation (Copernicus Climate Data Store, 2020). Additionally, we investigated how water surface occurrence and extent of the SLSW varied according to the size of water bodies. To accomplish this, the area (ha) of the identified water bodies was categorized into 5 classes, each reflecting a maximum number of associated water pixels. The classes ranged from 0.0064 to 0.0128 ha (up to 2 pixels), from 0.0128 to 0.0256 (up to 3 pixels), from 0.0256 to 0.0384 (up to 5 pixels), from 0.0384 to 0.064 (up to 10 pixels), and from 0.064 to 0.64 (up to 100 pixels).

Second, the agreement in surface water extent (objective *iii*; Figure 2g) was assessed through linear and quadratic regression modeling between SLSW (dependent variable) and LGSW (independent variable) models, under both wet (January-April 2021) and dry (June-September 2021) environmental conditions. Such a statistical procedure was separately repeated using the independent verification dataset (Figure 2d), and spatiotemporally coincident predictions (May 2022) from SLSW and LGSW models. For each analysis, the distribution values were compared as density plots, and linear and quadratic regressions were performed. The coefficient of determination (R^2^) was used as a measure of relatedness (Peña-Luque et al., 2021), and the slope (B1) was calculated using standardized major axis regression (Warton and Weber, 2002).

The statistical analysis and data synthesis were conducted in the R environment (v.4.2.0; R Core Team, 2021), using the following packages: ‘*blockCV*’ (v.2.1.4; Valavi et al., 2018), ‘*randomForest*’ (v.4.7-1.1; Liaw and Wiener, 2002), ‘*pdp*’ (v.0.8.0; Greemwell, 2017), and ‘*rgeos*’ (v.0.6-2; Bivan et al., 2017).

## 3. Results

### 3.1. Surface water occurrence inferred from SLSW

The analysis of surface water occurrence performance indicated a relatively low OOB error rate (Table 2), with a similar score (*μ*≈26%) between the water and non-water classifiers. The specificity (true negative rate) metric demonstrated better performance than sensitivity (true positive rate), as indicated in Table 2. Similar results were observed for Accuracy, AUC, and F1 metrics. The MCC metric, which is a more conservative measure, indicates a good level of agreement (MC=0.45) (Table 2). Also, the metric scores from RF models including Sentinel-1&2 metrics were comparatively similar to scores solely including Sentinel-2 metrics, hence with no remarkable support by the Sentinel-1 data in terms of water surface classification accuracy (Table SM3).

**Table2.**
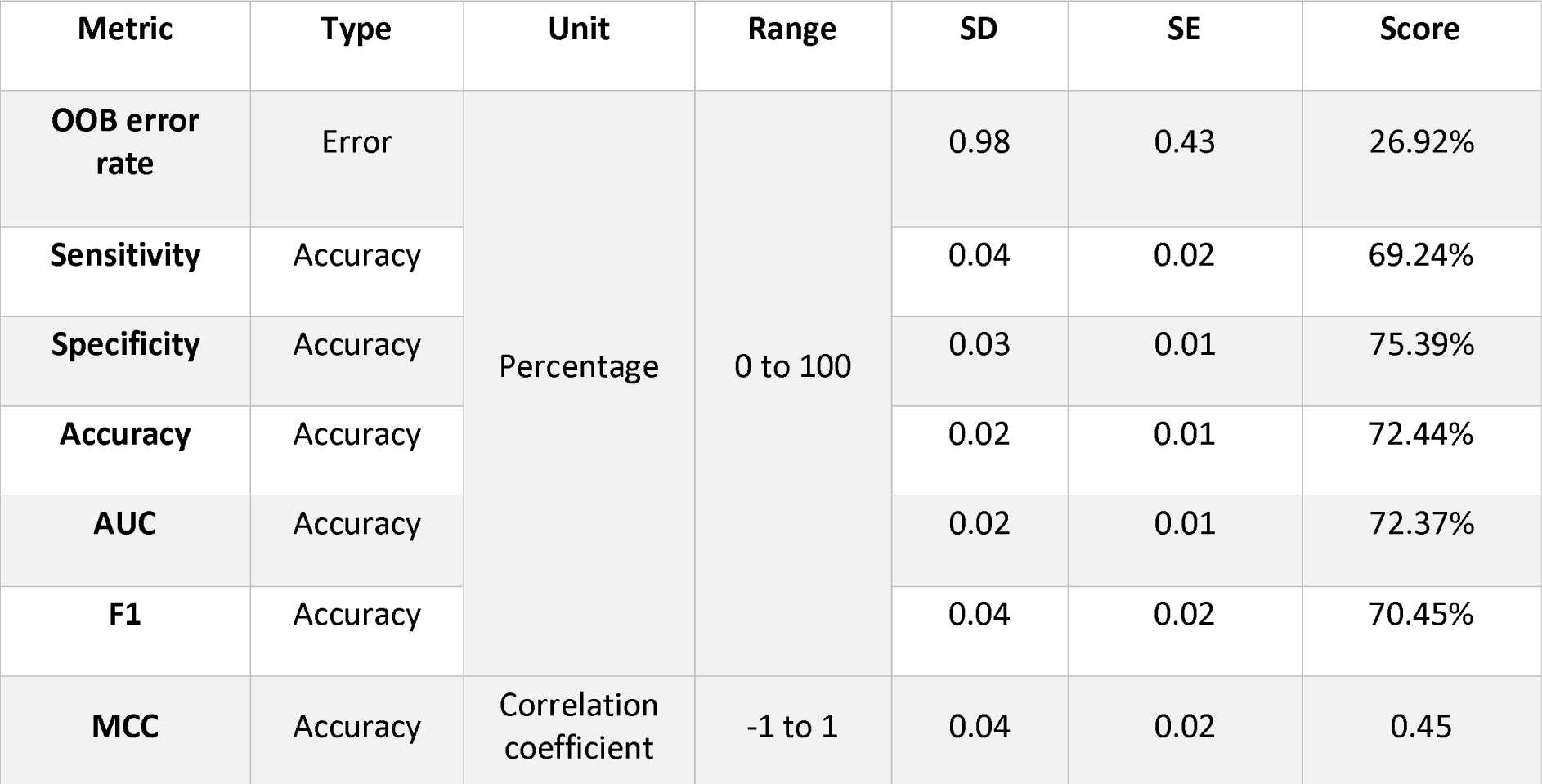
– Metrics scores (OOB (out-of-bag) error rate, Sensitivity, Specificity, Accuracy, AUC (Area Under the Curve), F1, and MCC (Matthews’ Correlation Coefficient)) defining RF model performances for surface water occurrence classification. Scores are coupled with standard deviation (SD) and standard error (SE), and averaged based on geographically distinct 5-fold RF cross-validations.

The results of RF analysis revealed similar rankings between variables that best explained water classification (water and non-water classes) (Figure 3a), and the variables that explained surface water occurrence (Figure 3b), according to the Gini index. More importantly, the importance cutoff indicated both Sentinel-1&2 data were significantly influential in explaining surface water occurrence (Figure 3b). Here, the most influential variables were the vegetation index NDI45, followed by the SWIR1, BLUE, and NIR1 reflectance bands. Second-order textural index GLCM-C and the SAR index polarization ratio VHVVR were also identified as influential variables in explaining surface water occurrence, as depicted in Figure 3b.

**Figure 3.**
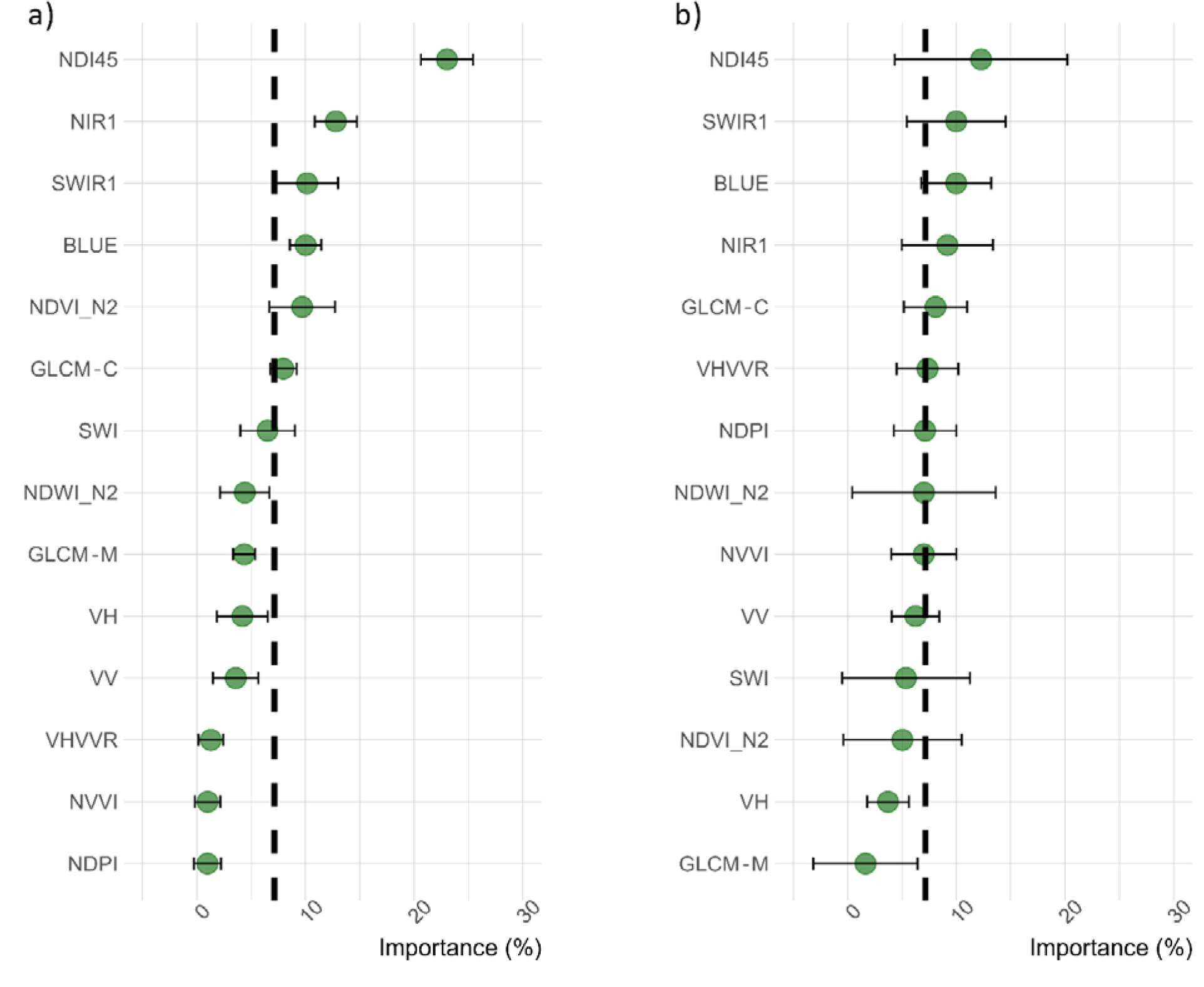
–Variable importance for a) water classification (water and non-water classes) and for b) surface water occurrence is expressed in percentage and averaged from the 5-fold cross-validations, where the mean is depicted by green dots, and error bars in black indicating the SD.

The probability of surface water occurrence improved with increasing NDI45 index values, reaching a plateau at around 0.25 (up to ∼0.5), as shown in Figure 4. Conversely, for SWIR1 and NIR1 reflectance bands, there were less steep curves with negative probabilities observed at the beginning of the bands’ reflectance value increase, corresponding to ∼0.2, while for the BLUE band, the negative probability was observed at around 0.05 (Figure 4). For the GLCM-C and VHVVR variables, highly steep curves (respectively positive and negative) were identified, with the former indicating a low probability at ∼0, but promptly reaching a plateau at ∼5 (up to ∼30), while for VHVVR, a higher probability was observed from ∼-100 to ∼0 backscattering values, followed by an abrupt negative response (Figure 4).

**Figure 4.**
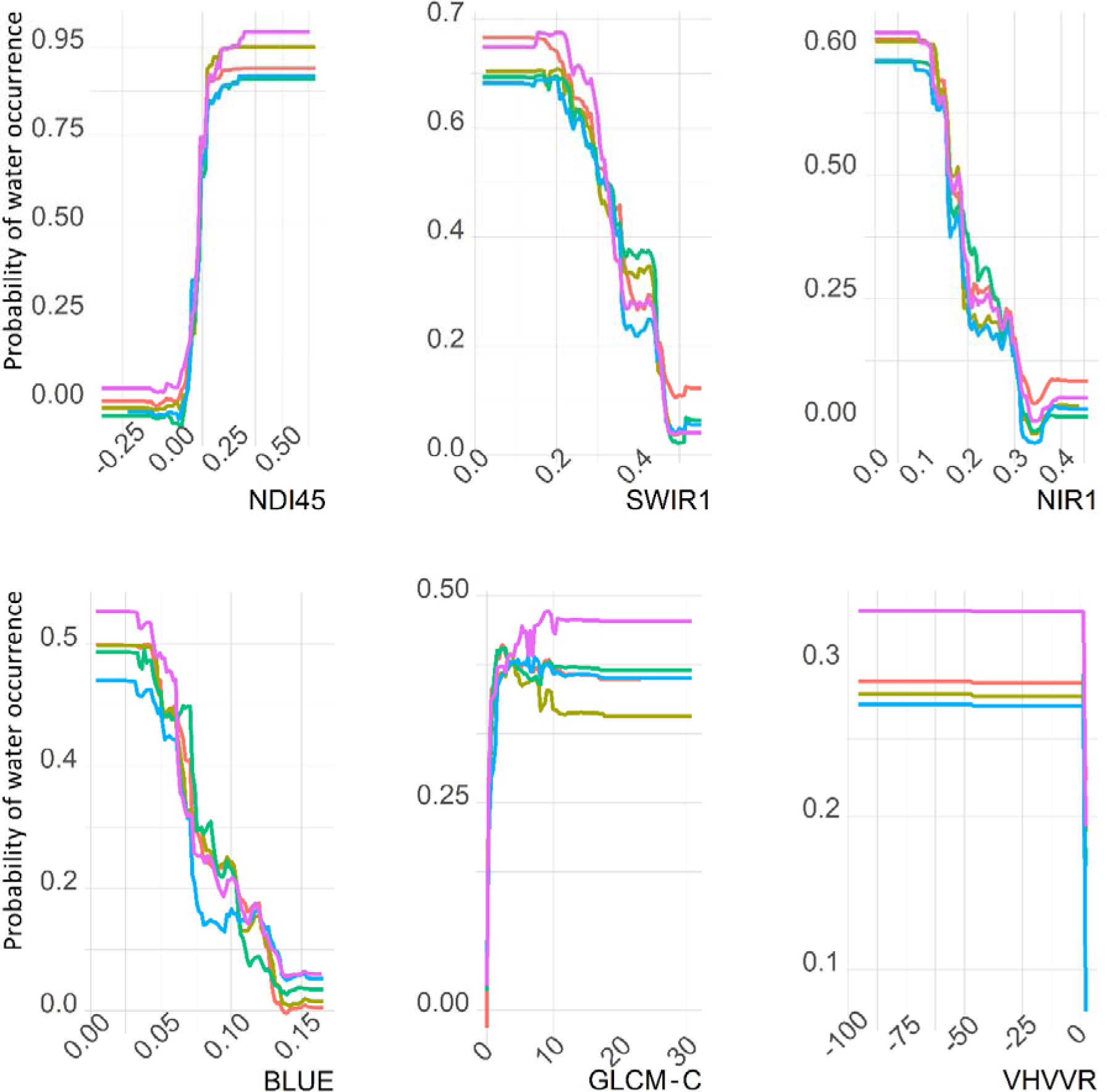
– Partial dependence plots of the most important variables explaining surface water occurrence. For each variable, several curves are depicted, with each color reflecting a fold of the stratified RF cross-validations. Additional plots of the remaining variables can be found in Supplementary Materials (Figure SM4).

### 3.2. Intra- and Inter-annual surface water trends inferred from SLSW

The intra- and inter-year trends in surface water attributes (occurrence and extent), from sampled water bodies, were estimated from SLSW. In the three regions combined, a clear seasonality pattern emerged, characterized by the magnitude of surface water changes. This reflected higher median values in surface water extent (_∼_60%) during autumn (November) to spring (late April), followed by a pronounced drop (_∼_-40%) during summer (Figure 5a). The minimum surface water extent conditions (_∼_20% to 10%) were observed for the SLSW from late July to early September, followed by an extensive recovery in November (Figure 5a). Both 2020 and 2021 exhibited similar seasonal variability, with high agreement during wetter conditions (November-April); yet, a prolonged anomalous period of water deficit conditions was detected from SLSW in 2020 between September to early October, where it can be observed slower recovery capabilities compared to 2021 (Figure 5a). It can be also observed that both water surface occurrence and extent prediction followed the spatiotemporal trends of the meteorological parameters derived from the independent E-OBS gridded data (SM5).

**Figure 5.**
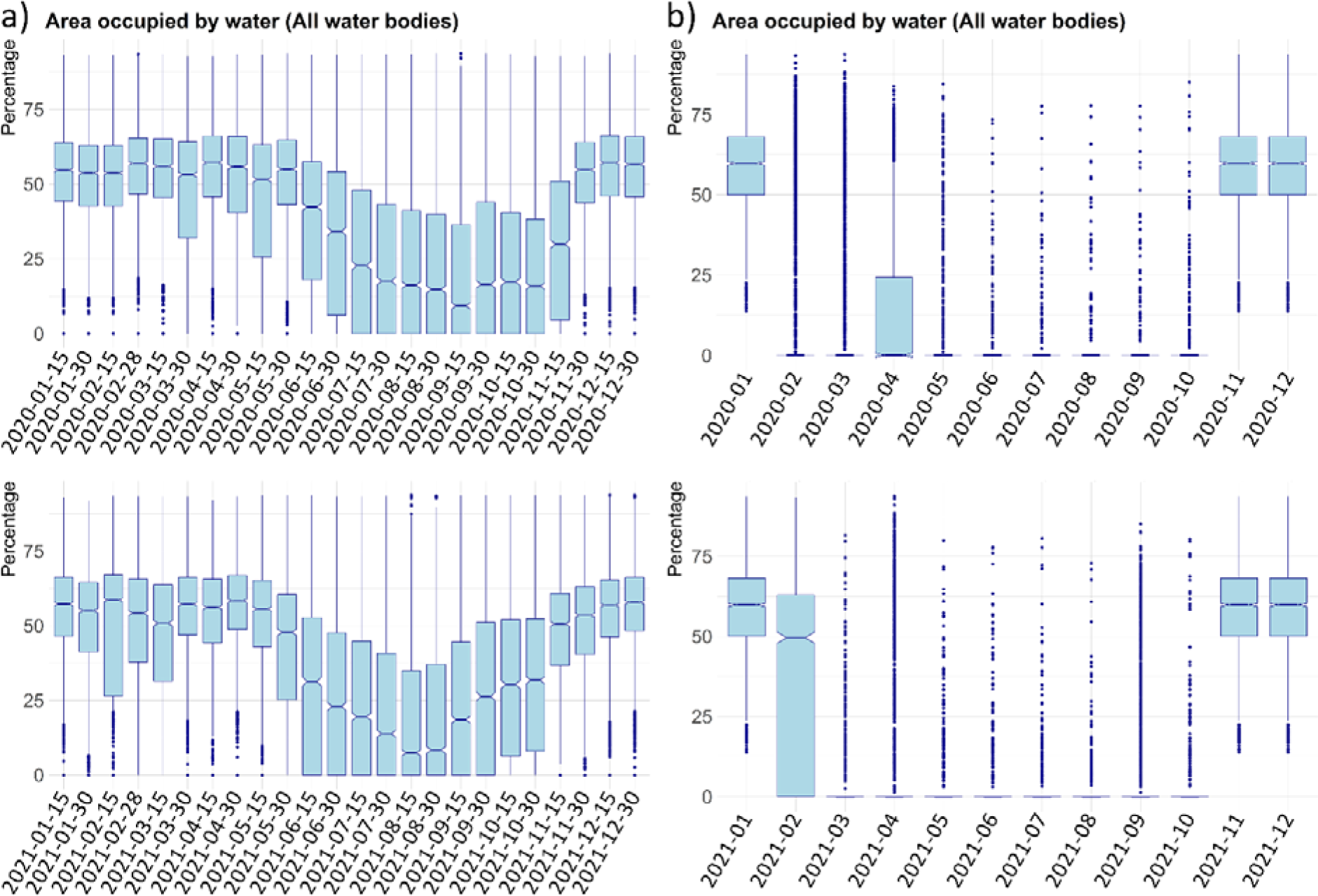
– Dynamic surface water extent, considering the three semiarid regions collectively. The time series are produced for the SLSW (a) and LGSW (b) models with a time interval of 15 days and 30 days, respectively. For each date, the range of values illustrates the surface water percentage in water bodies, with dots depicting the outliers. The time series of associated products were reproduced for the years 2020 (upper panels) and 2021 (lower panels).

When analyzed individually, the three regions exhibited distinct seasonal variability, with clear divergences observed when comparing intra- and inter-annual time series forecasting among the regions (Figure 6a). The region that held the strongest observed seasonal variability was SP1, followed by PT1 and then PT2, which presented the lowest intra-annual variability (Figure 6a). In terms of inter-annual seasonal variability, the three areas displayed similar trends during wetter conditions (November-April) (Figure 6a). However, during dryer conditions (summer), a larger deficit was observed in the time series, with a lack of recovery in dynamic surface water extent during 2020 compared to 2021, especially in PT1 and SP1 regions, yet with seemingly no difference for PT2 (Figure 6a).

**Figure 6.**
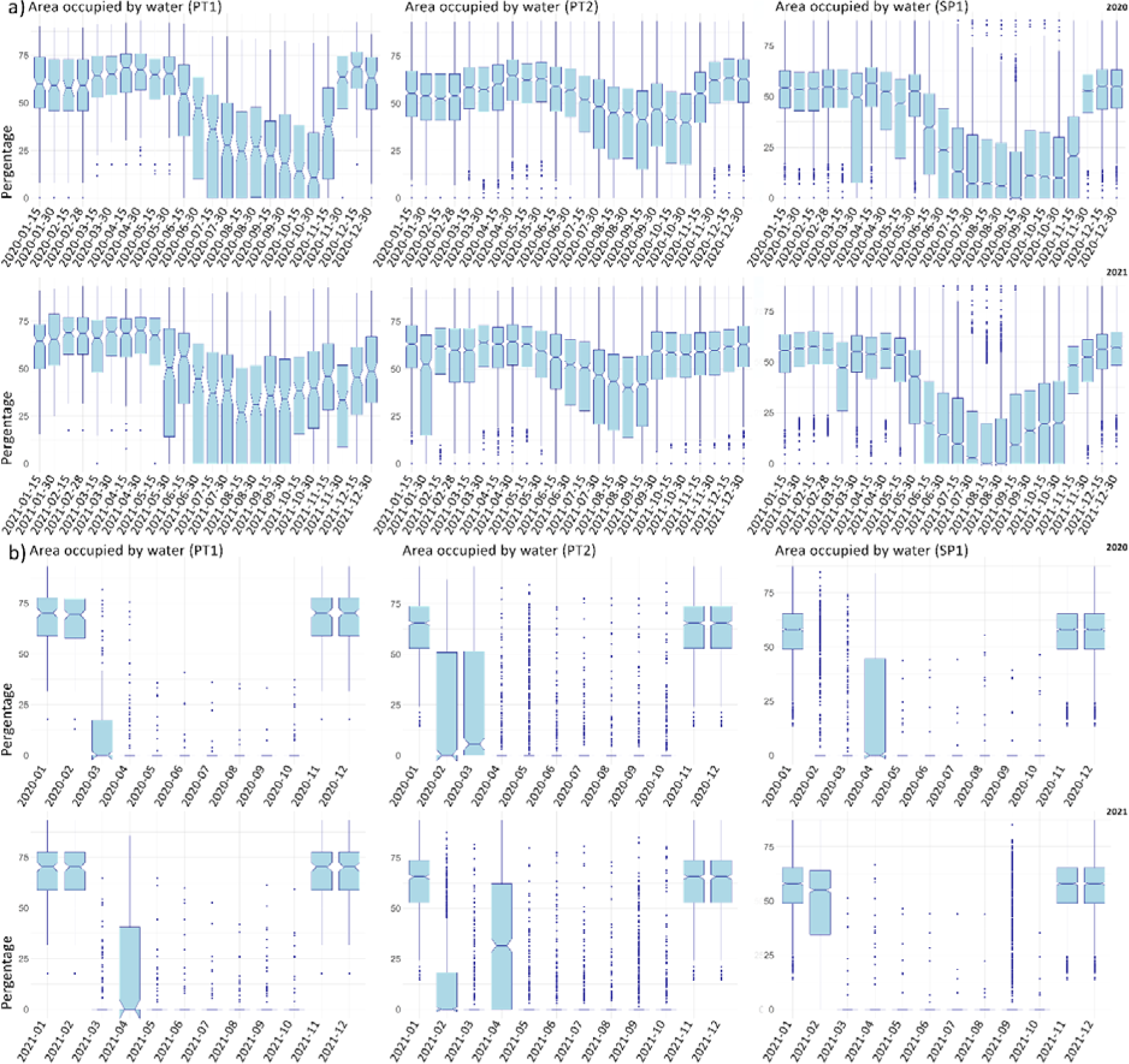
–Dynamic surface water extent, considering the three semiarid regions (PT1, PT2, and SP1) separately. For each date, the range of values illustrates the surface water percentage in water bodies, with dots depicting the outliers. The time series are produced for the SLSW (a) and LGSW (b) models with an interval of 15 days and 30 days, respectively. The time series of associated products were reproduced for the year 2020 and 2021.

The SLSW time series forecasting of dynamic surface water occurrence exhibited similar trends to those observed for water extent, both when considering the three regions as a whole (Figure SM6) and when analyzing them separately (Figure SM7). Diversely, when separating water bodies into size classes, both SLSW surface water occurrence and extent values demonstrated an upward trend coinciding with the enlargement of water body size in both 2020 and 2021 (Figure 7 and Figure SM8). Furthermore, the SLSW revealed a constant discrepancy between classes throughout the wet season (January-April), which became more pronounced during the dry season (June-September), particularly for the first class (0.0064-0.0128 ha) (Figure 7 and Figure SM8). Again, from mid-September to mid-October of 2020, anomalous extended periods of reduced surface water extent were evidenced compared to 2021, notably for the first class (Figure 7). Such temporary water deficit in this period coincided with higher temperature (mean and maximum), lower humidity and precipitation amount, and markedly lower values in surface shortwave downwelling radiation, compared to 2021 (SM5).

**Figure 7.**
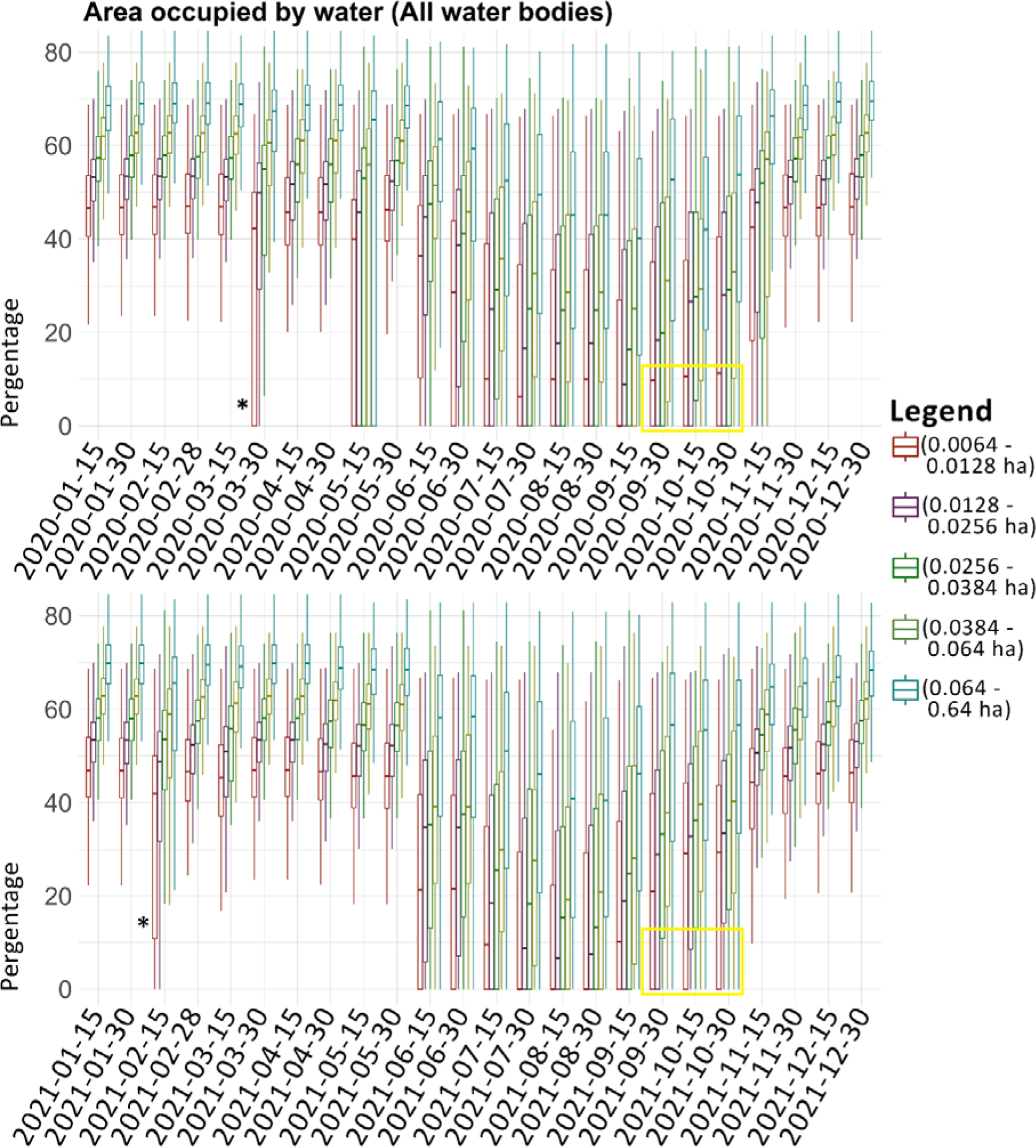
– Dynamic surface water extent (surface water percentage), where for each date, the range of values illustrates the variations according to the 5 size classes (0.0064-0.0128, 2 pixels; 0.0128-0.0256, 3 pixels; 0.0256-0.0384, 5 pixels; 0.0384-0.064, 10 pixels; 0.064-0.64, 100 pixels). The time series forecasting represents the SLSW product, and were reproduced for the years 2020 (upper panels) and 2021 (lower panels). The yellow box highlights the prolonged low surface water extent values (median value) in 2020, compared to 2021. The * symbols depict where boxplots indicate completeness problems during the wet period (January-April) (see Figure SM8 for further details).

### 3.3. SLSW and LGSW surface water inferences

#### 3.3.1 Archives completeness

The results for dynamic surface water extent trends were synthesized for SLSW and LGSW. Overall, complete archives emerged for SLSW contrary to LGSW for the three regions (Figures 5a, 5b, 6a, 6b). In fact, the LGSW time series forecasting repeatedly reported a lack of completeness (median values equal to 0), corresponding to an underestimation magnitude of approximately 3 times (*μ*≈70.5%) than SLSW, when considering all three regions together (Figure 5b). Nonetheless, a pronounced underestimation was also observed separately for PT1 and PT2 (*μ*≈66.6%), and SP1 (*μ*≈70.5%) (Figure 6b). Additionally, it was generally observed that the LGSW exhibited a higher underestimation of surface water under drier conditions (Figure 5b; Figure 6b). For the SLSW model, the time series forecasting consistently produced median surface water extent values above 0% (Figure 5a; Figure 6a). Yet, a thorough examination according to water bodies size classes evidenced some biases related to archive completeness, particularly among the first class (0.0064-0.0128 ha) (Figure 7). In fact, during specific dates (late March 2020 and early February 2021), the interquartile range of boxplots encompassed much lower values (∼10% for 2021 and 0% for 2020) in surface water extent compared to other dates within the wet season (Figure 7). Likewise, archive completeness issues also emerged on these dates for surface water occurrence, with classes displaying ∼20% decreases compared to other dates in the wet season (see Figure SM8). Furthermore, by taking the year 2021 as an example, the surface water occurrence mapping process showed more detailed information across the three regions (Figure 8a), and as expected, a higher representativeness of surface water occurrence pixels using SLSW rather than LGSW, in specific during drier periods (e.g., August) compared to wetter periods (e.g., January) (Figure 8b). The cumulative surface water occurrence associated with the SLSW is also depicted, highlighting pixels subject to higher and lower seasonal variability (seasonality) (Figure 8c).

**Figure 8.**
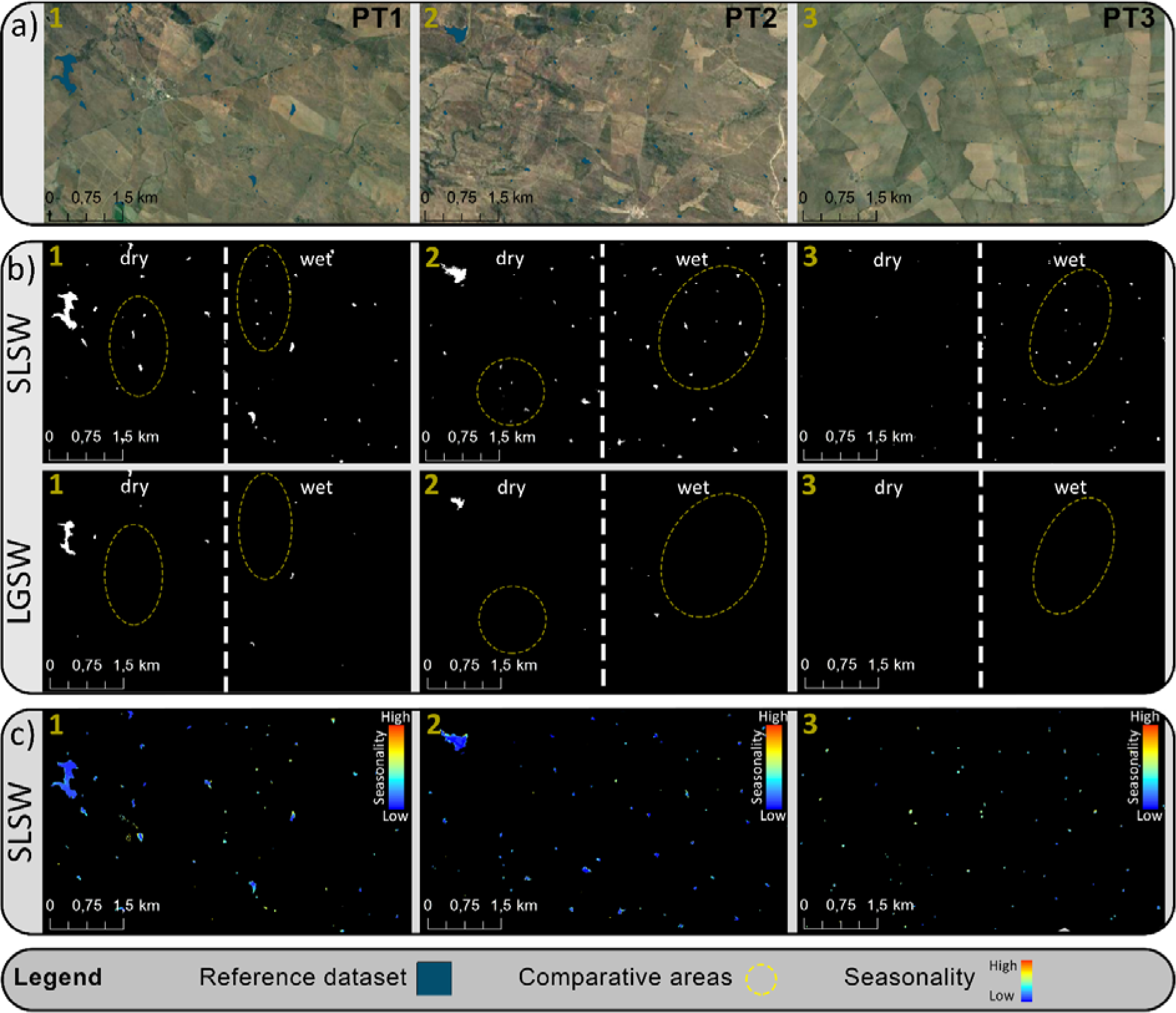
–Surface water distribution in the magnified areas (a), where numbers indicate their representative regions depicted in Figure1. The magnified areas (see Fig. 1) were comparable concerning surface water occurrence predictions (in white) by SLSW (upper panels) and LGSW (middle panels) (b), jointly with SLSW-derived seasonality (lower panels) (c). Upper and middle panels were bisected by a vertical (white) dashed line, to represent dry (August) and wet conditions (January). Ellipses and circles are also depicted in magnified areas highlighting clustered water bodies, to emphasize prediction comparisons between SLSW and LGSW.

Further results on surface water occurrence time series (SLSW and LGSW) can be found in Supplementary Materials (Figure SM6-SM9).

#### 3.3.2 Comparison between water bodies detection methods and verification dataset

When comparing regression models using SLSW as a function of LGSW for surface water extent, higher correlations (relatedness) were observed during the wet period (Figure 9a) compared to the dry period (Figure 9b), as evidenced by the linear and quadratic coefficients of determination (R^2^), and the slope (B1) of the regression. Yet, the correlations between both products are still low, with R^2^ values ranging from 0.05 (dry) to 0.38 (wet). The density plots revealed that SLSW overestimated surface water extent when compared with LGSW during both dry and wet periods, with a median difference of _∼_25% (Figure 9a, 9b). The magnitude of overestimation by SLSW was similar when compared with the verification data (_∼_25%) (Figure 9c). However, when SLSW and LGSW were separately compared with the verification data, SLSW exhibited better fit and performed well according to the three metrics (e.g., R^2^ = 0.66), while LGSW showed poor regression scores (Figure 9d). Additionally, the median difference from the density plots between LGSW and the verification data was markedly high, resulting in _∼_45% (Figure 9d), and evidencing an underestimation in surface water extent stronger than the overestimation observed with SLSW (Figure 9c).

**Figure 9.**
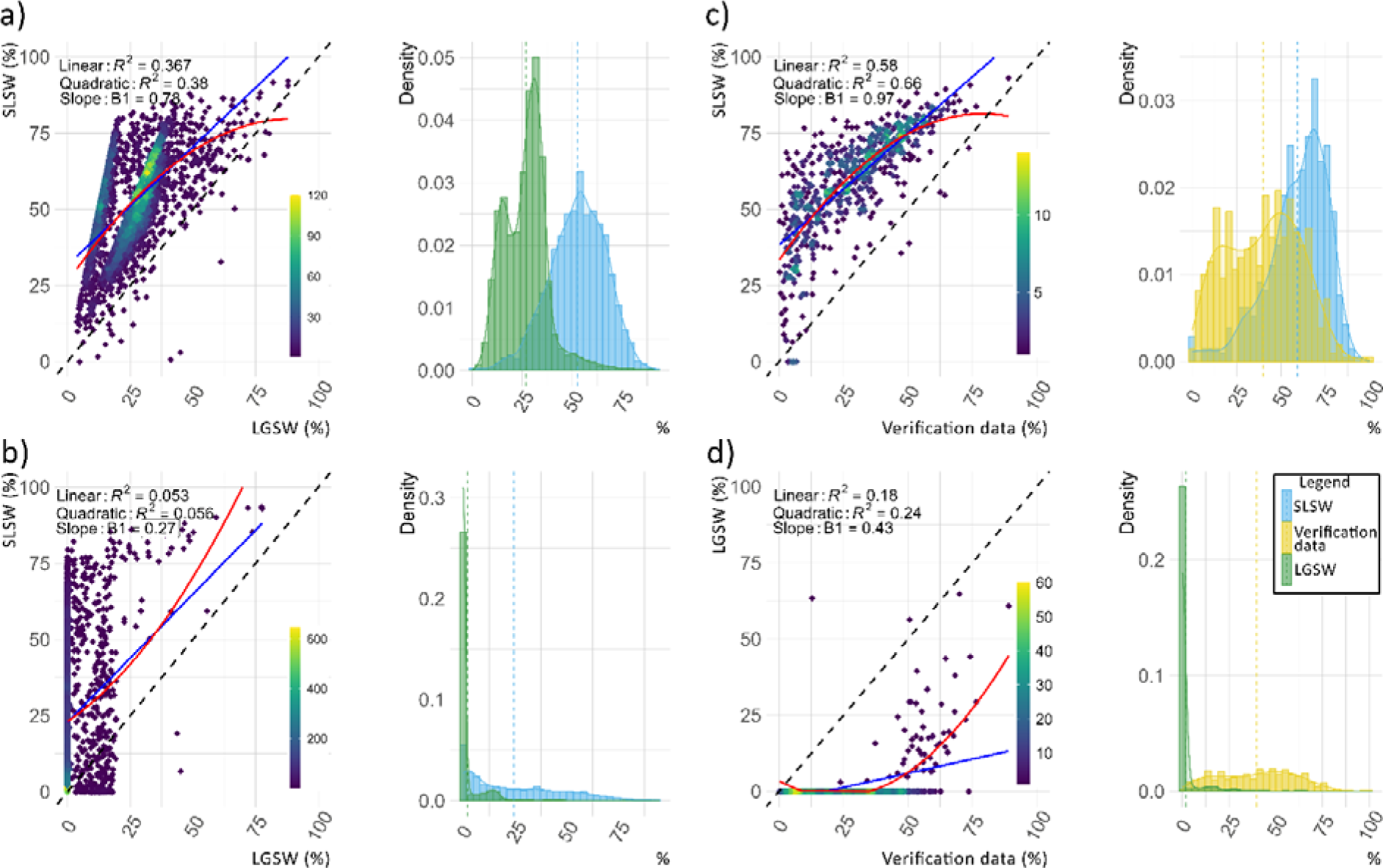
–Density scatterplots are depicted with the linear (blue line) and quadratic (red line) regressions between SLSW and LGSW concerning surface water extent (surface water percentage in water bodies) under both wet and dry conditions (a and b). Furthermore, independent comparisons of SLSW and LGSW with the verification dataset are presented in separate panels (c and d). In all panels, each graph point is a representation of a distinct water body. Density plots are coupled with each regression, where dashed lines represent the median value of estimated surface water extent.

## 4. Discussion

Our study proposes a multi-temporal multi-sensor data fusion method to bridge surface water catalog discontinuities, enhancing the characteristics delineation and mapping of small seasonal water bodies in semiarid environments. Results shed light on the prospective benefits of combining Sentinel-1 and Sentinel-2 data for surface water detection and quantification, and the value of time series to infer intra- and inter-year surface water trends in different areas. The reliability of our findings was supported by the comparative assessment between local and global products in terms of prediction completeness and agreement, focusing on wet and dry periods, and throughout an independent verification dataset.

### 4.1. Surface water occurrence inference

The NDI45 emerged as the most influential variable in explaining the potential occurrence of surface water. The strong relationship between NDI45 and chlorophyll content, as well as its high sensitivity in describing vegetation biophysical properties, using the red edge wavelength interval, is noteworthy (Jiang et al., 2022; Delegido et al., 2011). It has also been found to be effective in describing environmental characteristics in wetlands and semiarid regions (Jiang et al., 2022; Valerio et al., 2020). In contrast, previous research focused on water classification found lower NDI45 importance scores and higher scores for water-related indices with NDWI and MNDWI, which are possibly more suitable for distinguishing larger water areas (limnetic zone) (Jiang et al., 2022; Radoux et al., 2016), and clearer waters (Sun et al., 2012). The importance of NDI45 relies on its ability to discriminate between different types of vegetation, including both terrestrial and aquatic, and for the latter, minimum healthy conditions have been found at values ≈0.1 in a nearby geographic region (Soria et al., 2022). This finding aligns with the inflection point found in the NDI45 response curve, supporting the empirical evidence of aquatic plants in small water bodies such as ponds, which are often characterized by a diverse range of species (Casas et al., 2012; Céréghino et al., 2008). However, further ecological and conservation research is essential to gain a more evident understanding of the implications arising from this chlorophyll activity in relation to vegetation characteristics. In fact, it could be attributed to green waters with low biomass concentrations in well-represented littoral zones, or to eutrophication processes in various types of ponds, including those designated for livestock activities (Casas et al., 2012). The NIR and SWIR bands were also important as they explicitly detect water signals given the considerable absorption of electromagnetic radiation in the invisible wavelength area, whether non-water landscape attributes typically exhibit higher reflectance (Du et al., 2016; Mondejar and Tongco, 2019; Sun et al., 2012). Interestingly, a band from the visible spectrum, the BLUE band, also demonstrated high relevance, as indicated by the negative response curve, suggesting its association with water-related light absorbability. Yet, this band may be related to shallow areas due to its sensitivity to soil background reflection (Jiang et al., 2022), or to some degree of turbidity caused by suspended sediments (Sun et al., 2012). The horizontal structural GLMC_C variable indicated a rapid increase in the probability of surface water occurrence for areas with extremely contrasting values in photosynthetic activity. This suggests that the lowest and highest photosynthetic values corresponded to central water areas and vegetative buffer zones, respectively. This metric successfully captured the landscape ecological properties of water bodies, which may be beneficial in classification, extraction, and segmentation workflows, as well for accurate mapping of water bodies in uncharted regions (Bioresita et al., 2019; Du et al., 2016; Peña-Luque et al., 2021).

The prime variables used in this study were based on spectral information from Sentinel-2 sensors and radar information from Sentinel-1 sensors, facilitating surface water occurrence inference. One of the key metrics utilized was the cross-polarization ratioed SAR metric VHVVR (also known as VH/VV, or VHrVV). The backscattering response is influenced by the synergistic effects of the structure (e.g., volume scatters from vegetation), and the dielectric field of the surfaces, determining the resulting amount of energy received by the sensor (Vreugdenhil et al., 2020). Here, surface waters were mainly associated with very low backscatter values, which was an expected result because flat and open surface waters typically exhibit lower backscatter coefficients (e.g., Huang et al., 2018). A high probability of surface water occurrence was found with increased backscattered energy, ranging from highly smooth (lower backscatter coefficient) to more irregular surfaces (Tang et al., 2022). However, it is important to note that this range of values prior the inflection point, roughly around 0, may also include signals from wet soil and scattered vegetation, potentially including the detection of more irregularly inundated areas (Clement and Moore, 2018; Huang et al., 2018; Petropoulos et al., 2015). Interestingly, the VHVVR may hold the potential for conservation studies exclusively targeting temporary (semi-permanent) ponds, which can be more complex in terms of variant conditions (such as habitat, persistence, and pond bed; Sebastián-González and Green, 2014) compared to typical water bodies. As such, traditional approaches may face difficulties in comprehensively detecting and understanding such ponds, making the use of VHVVR particularly valuable in these cases.

### 4.2. SLSW and LGSW accuracy

#### 4.2.1 Intra and inter-annual surface water trends

The comparison between SLSW and LGSW for both 2020 and 2021 demonstrated a strong divergence in prediction capability for each region and across regions. Better completeness was observed from SLSW in both intra- and inter-annual surface water trends (occurrence and surface water extent). It is well-known that a major limitation of LGSW is the geographical and temporal discontinuities of the archives due to cloud cover conditions, especially for the detection of seasonal surface waters (Pekel et al., 2016). It is worth mentioning that during the wet season, recognized as the cloudiest period, specific dates were identified with missing information also in the SLWS time series forecasting. It is unlikely that this was caused by model misclassification, as both larger and smaller water bodies were affected likewise. Instead, the causes underlying this partial completeness can be found in regions where prolonged cloud cover occurs, resulting in a permanent pixel masking determined by the Quality Assessment (QA) bands. Considering this, it is also crucial to highlight that the amount of missing information was found to be negligible, meaning that such cloud persistence was circumscribed to local areas, and did not compromise the overall completeness of the archives at those dates. Hence, in line with previous research (Peña-Luque et al., 2021), the selected 15-day time windows appeared to be a reasonable compromise for ensuring the extraction of environmental data with a high spatial match, while maintaining reasonable temporal alignment. So, in short, the present temporospatial data fusion technique applied in semiarid environments effectively preserved high-frequency time series, providing sufficient information for gap filling procedures in semi-arid regions. Still, it’s important to acknowledge that LGSW holds a noteworthy benefit in handling research tasks that require EOS data spanning decades.

The SLWS corroborated high seasonal variability within the three areas, which can likely be explained by the temperature-related variations in evaporation and evapotranspiration rates, but also by the precipitation amount and relative humidity, which may vary considerably according to E-OBS trends. Among the three areas, the Natural Park of Vale do Guadiana (PT2) exhibited the lowest variation, possibly attributed to the larger presence of deep-rooted vegetation facilitating infiltration capacity, and thus groundwater recharge. In the other areas, by contrast, higher surface water losses during summer can be attributed to two reasons. Firstly, higher hydroclimatic variability, influenced by temperature and energy stocks, that typically affects smaller water bodies (Rachid et al., 2022), such as in La Serena y Sierras Periféricas (SP1). Secondly, the removal of water for human-related activities (irrigation, livestock, etc.) could also contribute to the decrease in water extent, in particular when resources from large neighbor reservoirs may not always be disposable, such as in Castro Verde (PT1). These findings reveal a recurrent pattern in surface water fluctuations, suggesting periodic changes in water pond levels, with water bodies conventionally filling from autumn to early winter (Gómez-Rodríguez et al., 2009). Nonetheless, the SLWS analysis indicated that 2020 experienced drier conditions than 2021, concomitantly with E-OBS data showing higher temperatures, lower humidity, and precipitation rates in 2020. Notably, the SLWS time series forecasting enabled the examination of fine-scale differences between years, uncovering water deficit anomalies particularly evident in smaller water bodies (0.0064 to 0.0128 ha) during the transition from the late dry season to the early wet season, attributable to prolonged dry conditions. The intra-year variation during this specific period was further evidenced by the E-OBS trends, particularly concerning the highest amplitude rates found in surface shortwave downwelling radiation, indicating a considerably increase in ground radiation absorbability during 2020. This result support potential intensification in water bodies’ energy stock and evaporation rates, as elucidated previously (Rachid et al., 2022). It also represents a new aspect deserving attention in future research, as the noted consecutive anomalies are associable with drought events, in turn traditionally detected at coarser scales (Zou et al., 2018). Climatic anomalies deriving from climatic change are suspected to increase in these regions, with rising temperatures and drought events posing socio-economic concerns for sustainable water resource management (Jenkins and Warren, 2015). These conditions also present a conservation challenge, especially for endangered wildlife species (e.g., *Tetrax tetrax*; *Otis tarda*), as prohibitive climatic conditions and impaired utilization of natural resources can have serious consequences during key phenological phases (Silva et al., 2015).

#### 4.2.2 Surface water inference reliability

In both dry and wet seasons, there was minimal agreement between SLSW and LGSW in terms of identified surface water extent. During dry periods, the surface water extent decreased dramatically, hindering the correlation between SLSW and LGSW, with the latter subjected to higher inaccuracy and underestimated surface water. This can be attributed to the inferior LGSW sensor grain (30m), which is a recognized limitation that needs to be addressed in future improvements of EOS-based approaches (Pekel et al., 2016). Therefore, it is crucial to emphasize the importance of the native Sentinel-1&2 resolution in the performance of SLSW models. Additionally, the higher spectral and radiometric resolution of SLSW, along with a wider wavelength spectrum, may have contributed to a better characterization of complex waters (Bangira et al., 2019), especially in water-stressed areas where frequent water shortages may interfere with sensors’ detection capabilities (Peña-Luque et al., 2021). The current findings also highlighted the importance of including more sensitive vegetation indices when describing complex waters, along with textural indices and the combination of different polarizations. Another possible reason for such contrast might be related to the archive’ discontinuities, although this is unlikely to be a significant factor as cloud cover is usually inconsistent during dry periods in many Mediterranean regions (Tulbure et al., 2022).

The limited ability of LGSW to accurately infer surface water extent in small ponds was further corroborated with the verification data, resulting in very poor agreement and a high underestimation of surface water extent. On the other hand, although SLSW exhibited considerably better performance, a slight tendency emerged in overestimating surface water extent. Nevertheless, this overestimation is likely a result of the algorithm’ classification *per se* rather than in an inadequately applied threshold. Multiple thresholds were combined with a machine learning technique to improve the discrimination of complex waters (e.g., Bangira et al., 2019), and the selected thresholding value was considered moderately precautionary in a similar study (moderate water confidence; Huang et al., 2018). To handle overestimations when extrapolating surface water occurrence in new areas, an interesting aspect deserving further investigation may lie in incorporating topographic predictors (e.g., topographic wetness index; Huang et al., 2018; Tang et al., 2022; Vanderhoof et al. 2023). In our methodology, polygon centroids were employed to ensure that the model could effectively extrapolate its predictions, irrespective of the size of the water bodies involved. Yet, it is important to note that caution is necessary when incorporating large reservoirs with purer waters, as it may require careful scrutiny of predictions. For studies interested in surface water extent, given the variegated nature of ponds, a prudent recommendation is to predefine pond boundaries before implementing extrapolations, which may otherwise result more time-consuming.

### 4.3. Limitations and opportunities

#### 4.3.1 SLSW limitations

When targeting permanent waters in relatively large reservoirs, previous academic work has usually achieved high accuracy scores, ranging from 70% to over 90% (Bioresita et al., 2019; Jiang et al., 2022; Zhou et al., 2017). Similar outcomes have also been evidenced for seasonal waters (Bangira et al., 2019; Huang et al., 2018). For small water bodies, recent studies have reported high accuracy in surface water occurrence and extent for sizes ranging from 0.5 ha (Wang et al., 2022) up to 5 ha (DeLancey et al., 2022; Peña-Luque et al., 2021). To date, there is a scarcity of research specifically addressing very small water bodies (<0.5 ha), and the few existing studies focusing on this range of sizes (0-0.5 ha) have consistently demonstrated suboptimal performance, especially when employing publicly available EOS data (Cordeiro et al., 2021). Virtually, no studies exist focused on mapping small water bodies comparable to our study (*Mdn*≈0.031 ha), while also investigating their complex seasonal dynamics. It is worth mentioning, however, that a slight underestimation of surface water occurrence, attributable to prediction uncertainties, may have occurred for water bodies within the smaller size-class (0.0064 to 0.0128 ha). This inaccuracy, despite being marginal, may have arisen from factors such as sensor resolution, algorithm sensitivity, or other pertinent technical considerations resulting for instance from pond destruction or displacement. Besides, one key limitation in our research is that those water bodies exhibit a diminished ability to reach full water surface extent capacity. This may be due to the increased susceptibility of shores to be misconstrued with areas affected by flood events. Flood-prone shores within small water bodies may attain full capacity only intermittently during flooding events, being typically ephemeral in semiarid environments (Tulbure et al., 2022). An alternative explanation for the discernible value separation across size classes lies in the 8-meter buffer analyses implemented on water bodies before computing surface water extent, notwithstanding this operation was essential to address errors originating from pixels that were either partially or largely situated outside water bodies. Despite some limitations, the metrics scores, the reproduced time series, and the validation can be considered satisfactory, given the challenging task of detecting water surfaces that are prone to alterations (Wang et al., 2022), especially in very small ponds with intermittent water levels and underlying seasonal effects, such as fluctuations in turbidity, background soil reflectance, and photosynthetic activity (Bangira et al., 2019; Das et al., 2021).

To soften biases more effectively, Sun et al. (2012) have suggested separating water bodies into three categories: clear, green, and turbid water. Still, the detected error arising from surface water occurrence analyses might be due to mixed pixels, which frequently provide the sole information for describing environmental characteristics, and therefore cannot be ignored. In this context, the literature widely acknowledges the efficacy of employing sub-pixel approaches (Radoux et al., 2016; Ozesmi & Bauer, 2002). Caution is needed, however, on this issue as multiple distinct classifications to derive spectral signatures can significantly increase computation time, which is especially relevant when forecasting at high frequency. A possible alternative solution to reduce biases involves omitting the buffer analysis and reclassifying any surface water extent value exceeding 100% to this maximum value. Lastly, while it is believed that a complete correction of false detections is unrealistic (Peña-Luque et al., 2021), reducing surface water extent overestimation requires algorithm improvements to address pixel misclassification issues, such as incorporating deep learning techniques (Mayer et al., 2021). However, deep learning methods demand more extensive datasets, often exceeding 10,000 samples, and still have consistent barriers in interpretability (see Fan et al., 2021).

#### 4.3.2 Applications in Ecology

Water availability is critical for the persistence of plant and wildlife populations, and the development of human activities, particularly in water-stressed and drought-prone regions. The present framework can be of special interest for a range of research and application fields in semiarid environments. It can contribute to the restoration/conservation of small wetland areas, increasingly encouraged by initiatives such as the Ramsar Convention (De Meester et al., 2005; Gxokwe et al., 2022; Slagter et al., 2020). The predominance of Sentinel-2 data in explaining water surface occurrence revealed that the inclusion of Sentinel-1 data may provide some gains in open water scenarios, for instance when targeting specific ponds (i.e., temporary ponds). It is also reasonable to hypothesize that Sentinel-1 data may prove more significant in more structured wetland environments, notably in vegetated waters (*sensu* Vanderhoof et al. 2023), where a variegated data on soil and vegetation characteristics is often required (Shen et al., 2019). In light of this, wetland habitat classification via remote sensing remains an open topic, enclosing various challenges that need careful addressing (Ozesmi & Bauer, 2002), especially dynamic environments (Tulbure et al., 2022). Ecological applications of our approach and framework can, therefore, be extended to small-sized wetlands influenced by seasonal variability and intermittent inundations, thus aiding in the recognition of highly overlooked wetland typologies (e.g., Semeniuk & Semeniuk, 1995). Such cases in fact require tailored approaches with detailed spatiotemporal information and including long periods for proper monitoring. The extensive lifespan and frequent site revisitation leveraging public satellites may prove competitive, even compared to commercial ones, including airborne sensor systems. This is because while such systems are acknowledged for their superior spatial and spectral resolution (Prošek et al., 2020; Sawaya et al., 2003), they often incur high costs and are conventionally subjected to limitations in terms of temporal resolution and swath, and when considering long-term studies. Commercial datasets may indeed be subject to restricted availability in online repositories (e.g., GEE). The highly detailed resolution of commercial data exacerbates the constraints, making metrics calculations prohibitively time-consuming when conducted outside cloud-computing platforms, especially when dealing with high-frequency time series. Long-term environmental monitoring provided by public sensors may also prove useful in the protection and restoration of temporary ponds (Céréghino et al., 2008; De Meester et al., 2005). This is relevant considering the role of small-sized ponds as habitat for conservation-priority species (Zamora-Marín et al., 2024), along with the fact that those ponds are often neglected or utilized unsustainably (Gleick, 1998; Olmo et al., 2022). Interestingly, the observation of inter-annual water deficit anomalies can provide insights into emerging threats (see Reid et al., 2018) and their implications, which are otherwise difficult to detect on pond hydroperiods, on which local biodiversity depends (De Meester et al., 2005). By enabling the detection of open surface water, this framework also provides a step forward in measuring the spatiotemporal arrangement and effective connectivity of small-sized ponds, useful for prioritizing their protection and wildlife management (Sebastián-González and Green, 2014). The consideration of forecasting at high frequency also supports the shifting paradigm from a static unrealistic landscape viewpoint to a dynamic landscape connectivity approach (Martensen et al., 2017; Zeller et al., 2020). This may facilitate better planning solutions to guarantee population viability for different taxonomic groups of plants and wildlife species, based on connectivity constraints imposed by water presence and species dispersal potential (Herceg-Szórádi and Csergő, 2023; Tucker et al., 2018; Vasudev et al., 2015).

#### 4.3.3 Applications for water sustainable use

Sustainable water resources management and development arouse interest not only in water detection, but also in water quality assessment (Gholizadeh et al., 2016; Ritchie et al., 2003). Ponds are vulnerable to direct or indirect contamination from pollutants and overexploitation in many regions, deepening the challenge of achieving equilibrium between rural competitiveness and ecological sustainability (Poláková et al., 2011; Reid et al., 2018). In the face of a changing climate and a growing human population, adaptive strategies are necessary, particularly concerning water resources in semiarid regions (Gašparović and Singh, 2022; Scholes 2020; Wickens, 1998). Remote sensing applications, as demonstrated in this study, may be useful for future research lines expanding the scope to quality assessments in open water bodies and across wide areas, providing new opportunities to address environmental challenges and aligning with SDGs (Bhaduri et al., 2016). A pressing need regards also novel epistemological frameworks to promote innovation in water resource monitoring and drought event detection, given the growing interest in multiple impactful areas (Anderson et al., 2023). Additionally, our modelling procedure could be beneficial for initiatives devoted to the reconstruction of hydrological cycles (Gleick, 1998), and to ameliorate renewable water resources in places without significant reservoirs, or to promote the planning of supplementary and alternative water bodies (Erwin, 2009; Morante-Carballo et al., 2022). In semiarid regions, often characterized by poor governance, such initiatives can be valuable for achieving policy objectives (i.e., food security and public health), and thus for uncovering the spatiotemporal distribution of social impacts arising from water scarcity, which is expected to increase significantly in Europe (Samaniego et al., 2018). Therefore, our approach may provide key information for improving water resource management protocols and advancing SDGs, fostering negotiations for conflict resolutions between water governance and environmental sustainability, involving institutions, stakeholders, policymakers, and local communities (Esgalhado et al., 2021; Jeffrey and Gearey, 2006; Veldkamp et al., 2015).

## 5. Conclusions

This research addresses contemporary gaps in water surface mapping, underscoring the benefits of employing high spatiotemporal resolution EOS imagery for empirical monitoring of small inland freshwater systems in semiarid regions. The utilization of this imagery reveals numerous advantages, particularly in quantifying hydrological dynamics and through continuous data analysis (Coops and Wulder, 2019). The robustness of the presented research was supported through accuracy metrics, model comparison with recent state-of-the-art research, and validations using independent verification data. Cautions were therefore raised towards traditional water surface products applied in this context, notably those derived from Landsat. Landsat-derived products seem better suited for larger and less complex water bodies due to their lower spatial and spectral resolution, whilst their mission longevity makes them suitable for long-term analysis, given the long-standing information spanning decades (Halabisky et al., 2016; Pekel et al., 2016). On the other hand, additional comparative assessments are suggested to validate the forthcoming Opera Dynamic Surface Water Extent (DSWE) product (Version 1), which is derived from harmonized Landsat and Sentinel-2 datasets (Mapper and Plus, 2023; Bato et al., 2023). This is also valid for the upcoming Landsat Next, known to be featured with high spectral resolution (https://www.usgs.gov/landsat-missions/landsat-next). Sentinel-1 and Sentinel-2 data offer a greater contribution to addressing water-related archive inconsistencies, particularly for very small-sized bodies, and the characterization of their complex dynamics, including the detection of consecutive anomalies. This is made possible by recent technological advancements in optical and radar sensors, providing high-quality and high-resolution information that is readily available in online repositories and can be processed using cloud computing power (e.g., GEE). The presented framework finally supports creative pathways for advancing land and water monitoring in a plethora of applications, bringing novel solutions to cross-boundary conservation and sustainability strategies (Radočaj et al.,2020; Wulder and Coops, 2014).

## Code availability

The GEE and R codes are available on GitHub at https://github.com/FrankVal/WatSurf.

## Author contributions

Francesco Valerio: Conceptualization, Data curation, Methodology, Software, Formal Analysis, Writing - Original Draft, Writing - Review & Editing, Visualization. Sérgio Godinho: Conceptualization, Methodology, Investigation, Validation, Supervision, Writing - Review & Editing, Visualization. Gonçalo Ferraz: Data curation, Writing - Review & Editing. Ricardo Pita: Supervision, Writing - Review & Editing. João Gameiro: Supervision, Writing - Review & Editing. Bruno Silva: Software, Validation, Writing - Review & Editing. Ana Teresa Marques: Methodology, Supervision, Writing - Review & Editing, Visualization. João Paulo Silva: Conceptualization, Data curation, Methodology, Writing - Review & Editing, Funding acquisition, Project administration.

## Acknowledgments

Work supported by National Funds through FCT-Fundação para a Ciência e a Tecnologia in the scope of the project UIDP/50027/2020. This work, including the scholarships of FV and ATM, was co-funded by the project NORTE-01-0246-FEDER-000063, supported by Norte Portugal Regional Operational Programme (NORTE2020), under the PORTUGAL 2020 Partnership Agreement, through the European Regional Development Fund (ERDF). FCT funded also BS through the grant SFRH/BD/137803/2018, while RP through the contract 2022.02878.CEECIND, and JPS through the contract DL57/2019/CP 1440/CT 0021. JG was supported by a post-doc scholarship (BIOPOLIS 2022-13). SG was funded by the FUEL-SAT project “Integration of multi-source satellite data for wildland fuel mapping: the role of remote sensing for an effective wildfire fuel management” from Foundation for Science and Technology (PCIF/GRF/0116/2019), and by National Funds through FCT under the Project UIDB/05183/2020.

## References

AEMET, IMP (2011) Atlas climático ibérico/Iberian climate atlas. Agencia Estatal de Meteorología, Ministerio de Medio Ambiente y Medio Rural y Marino, Instituto de Meteorologia de Portugal, Madrid.

Alvarez-Mozos, J., Villanueva, J., Arias, M., & Gonzalez-Audicana, M. (2021, July). Correlation between NDVI and Sentinel-1 derived features for maize. In 2021 IEEE International Geoscience and Remote Sensing Symposium IGARSS (pp. 6773–6776). IEEE. doi: 10.1109/IGARSS47720.2021.9554099

Anderson, K., Tooth, S., Kim, D., Resler, L. M., Schillereff, D., Williams, J. W., et al (2023). A horizon scan for novel and impactful areas of physical geography research in 2023 and beyond. Progress in Physical Geography: Earth and Environment, 03091333231217881. doi: 10.1177/03091333231217881

Millennium Ecosystem Assessment (2005). Ecosystems and human well-being: wetlands and water. World Resources Institute.

Attema, E., Davidson, M., Floury, N., Levrini, G., Rosich, B., Rommen, B., & Snoeij, P. (2008, June). Sentinel-1 ESA’s new European radar observatory. In 7th European conference on synthetic aperture radar (pp. 1-4). VDE.

Bangira, T., Alfieri, S. M., Menenti, M., & Van Niekerk, A. (2019). Comparing thresholding with machine learning classifiers for mapping complex water. Remote Sensing, 11(11), 1351. doi: 10.3390/rs11111351

Balázs, B., Bíró, T., Dyke, G., Singh, S. K., & Szabó, S. (2018). Extracting water-related features using reflectance data and principal component analysis of Landsat images. Hydrological Sciences Journal, 63(2), 269–284. doi: 10.1080/02626667.2018.1425802

Baldi, P., Brunak, S., Chauvin, Y., Andersen, C. A., & Nielsen, H. (2000). Assessing the accuracy of prediction algorithms for classification: an overview. Bioinformatics, 16(5), 412–424. doi: 10.1093/bioinformatics/16.5.412

Bato, M. G., Devlin, K., Dhillon, R., Bonnema, M., Sangha, S., Niemoeller, S., et al (2023). *A first look at the OPERA Surface Water eXtent and Land Surface Disturbance products and their applications* (No. EGU23-10200). Copernicus Meetings.

Bhaduri, A., Bogardi, J., Siddiqi, A., Voigt, H., Vörösmarty, C., Pahl-Wostl, C., et al (2016). Achieving sustainable development goals from a water perspective. Frontiers in Environmental Science, 64. doi: 10.3389/fenvs.2016.00064

Bioresita, F., Puissant, A., Stumpf, A., & Malet, J. P. (2019). Fusion of Sentinel-1 and Sentinel-2 image time series for permanent and temporary surface water mapping. International Journal of Remote Sensing, 40(23), 9026–9049. doi: 10.1080/01431161.2019.1624869

Bijeesh, T. V., & Narasimhamurthy, K. N. (2020). Surface water detection and delineation using remote sensing images: A review of methods and algorithms. Sustainable Water Resources Management, 6, 1–23. doi: 10.1007/s40899-020-00425-4

Bivand, R., Rundel, C., & Pebesma, E. (2017). rgeos: interface to geometry engine-open source (GEOS). R package version 0.3-26.

Breiman, L. (2001). Random forests. Machine learning, 45, 5–32. doi: 10.1023/A:1010933404324

Cao, Y. G., Yan, L. J., & Zheng, Z. Z. (2008). Extraction of information on geology hazard from multi-polarization SAR images. *The International Achieves of the Photogrammetry*, Remote Sensing and Spatial Information Science, 37, 1529–1532.

Casas, J. J., Toja, J., Peñalver, P., Juan, M., León, D., Fuentes-Rodríguez, F., et al (2012). Farm ponds as potential complementary habitats to natural wetlands in a Mediterranean region. Wetlands, 32, 161–174. doi: 10.1007/s13157-011-0265-5

Chen, Z., & Zhao, S. (2022). Automatic monitoring of surface water dynamics using Sentinel-1 and Sentinel-2 data with Google Earth Engine. International Journal of Applied Earth Observation and Geoinformation, 113, 103010. doi: 10.1016/j.jag.2022.103010

Clement, M. A., Kilsby, C. G., & Moore, P. (2018). Multi-temporal synthetic aperture radar flood mapping using change detection. Journal of Flood Risk Management, 11(2), 152–168. doi: 10.1111/jfr3.12303

Cornes, R. C., van der Schrier, G., van den Besselaar, E. J., & Jones, P. D. (2018). An ensemble version of the E-OBS temperature and precipitation data sets. Journal of Geophysical Research: Atmospheres, 123(17), 9391-9409. doi: 10.1029/2017JD028200

Céréghino, R., Biggs, J., Oertli, B., & Declerck, S. (2008). The ecology of European ponds: defining the characteristics of a neglected freshwater habitat. Hydrobiologia, 597, 1–6. doi: 10.1007/s10750-007-9225-8

Cochran, W. G. (1977). Sampling techniques. John Wiley & Sons.

Coops, N. C., & Wulder, M. A. (2019). Breaking the Habit (at). Trends in Ecology & Evolution, 34(7), 585–587. doi: 10.1016/j.tree.2019.04.013

Copernicus Climate Change Service, Climate Data Store, (2020): E-OBS daily gridded meteorological data for Europe from 1950 to present derived from in-situ observations. Copernicus Climate Change Service (C3S) Climate Data Store (CDS). DOI: 10.24381/cds.151d3ec6 (Accessed on 07-12-2023)

Cordeiro, M. C., Martinez, J. M., & Peña-Luque, S. (2021). Automatic water detection from multidimensional hierarchical clustering for Sentinel-2 images and a comparison with Level 2A processors. Remote Sensing of Environment, 253, 112209. doi: 10.1016/j.rse.2020.112209

Das, N., Bhattacharjee, R., Choubey, A., Ohri, A., Dwivedi, S. B., & Gaur, S. (2021). Time series analysis of automated surface water extraction and thermal pattern variation over the Betwa river, India. Advances in Space Research, 68(4), 1761–1788. doi: 10.1016/j.asr.2021.04.020

DeLancey, E. R., Czekajlo, A., Boychuk, L., Gregory, F., Amani, M., Brisco, B., et al (2022). Creating a Detailed Wetland Inventory with Sentinel-2 Time-Series Data and Google Earth Engine in the Prairie Pothole Region of Canada. Remote Sensing, 14(14), 3401. doi: 10.3390/rs14143401

Delegido, J., Verrelst, J., Alonso, L., & Moreno, J. (2011). Evaluation of sentinel-2 red-edge bands for empirical estimation of green LAI and chlorophyll content. Sensors, 11(7), 7063–7081. doi: 10.3390/s110707063

De Meester, L., Declerck, S., Stoks, R., Louette, G., Van De Meutter, F., De Bie, T., et al (2005). Ponds and pools as model systems in conservation biology, ecology and evolutionary biology. Aquatic conservation: Marine and freshwater ecosystems, 15(6), 715–725. doi: 10.1002/aqc.748

Du, Y., Zhang, Y., Ling, F., Wang, Q., Li, W., & Li, X. (2016). Water bodies’ mapping from Sentinel-2 imagery with modified normalized difference water index at 10-m spatial resolution produced by sharpening the SWIR band. Remote Sensing, 8(4), 354. doi: 10.3390/rs8040354

Diffenbaugh, N. S., Pal, J. S., Giorgi, F., & Gao, X. (2007). Heat stress intensification in the Mediterranean climate change hotspot. Geophysical Research Letters, 34(11). doi: 10.1029/2007GL030000

Downing, J. A., Prairie, Y. T., Cole, J. J., Duarte, C. M., Tranvik, L. J., Striegl, R. G., et al (2006). The global abundance and size distribution of lakes, ponds, and impoundments. Limnology and oceanography, 51(5), 2388–2397. doi: 10.4319/lo.2006.51.5.2388

Drusch, M., Del Bello, U., Carlier, S., Colin, O., Fernandez, V., Gascon, F., et al (2012). Sentinel-2: ESA’s optical high-resolution mission for GMES operational services. Remote sensing of Environment, 120, 25–36. doi: 10.1016/j.rse.2011.11.026

Erwin, K. L. (2009). Wetlands and global climate change: the role of wetland restoration in a changing world. Wetlands Ecology and management, 17(1), 71–84. doi: 10.1007/s11273-008-9119-1

Esgalhado, C., Guimarães, M. H., Lardon, S., Debolini, M., Balzan, M. V., Gennai-Schott, S. C., et al (2021). Mediterranean land system dynamics and their underlying drivers: Stakeholder perception from multiple case studies. Landscape and Urban Planning, 213, 104134. doi: 10.1016/j.landurbplan.2021.104134

Fan, F. L., Xiong, J., Li, M., & Wang, G. (2021). On interpretability of artificial neural networks: A survey. IEEE Transactions on Radiation and Plasma Medical Sciences, 5(6), 741–760. doi: 10.1109/TRPMS.2021.3066428

Gao, B. C. (1996). NDWI—A normalized difference water index for remote sensing of vegetation liquid water from space. Remote sensing of environment, 58(3), 257–266. doi: 10.1016/S0034-4257(96)00067-3

Gašparović, M., & Singh, S. K. (2022). Urban surface water bodies mapping using the automatic k-means based approach and sentinel-2 imagery. Geocarto International, 2148757. doi: 10.1080/10106049.2022.2148757

Gleick, P. H. (1998). Water in crisis: paths to sustainable water use. Ecological applications, 8(3), 571–579. doi: https://www.jstor.org/stable/2641249

Google, Inc., Google Earth software, http://earth.google.com/ [last accessed on March 13, 2022].

Gorelick, N., Hancher, M., Dixon, M., Ilyushchenko, S., Thau, D., & Moore, R. (2017). Google Earth Engine: Planetary-scale geospatial analysis for everyone. R e m o t e s e n s i n g o, 2f02, E1n8-2v7.i rdooi: n m e n t 10.1016/j.rse.2017.06.031

Greenwell, B. M. (2017). pdp: an R Package for constructing partial dependence plots. R J., 9(1), 421.

Gholizadeh, M. H., Melesse, A. M., & Reddi, L. (2016). A comprehensive review on water quality parameters estimation using remote sensing techniques. Sensors, 16(8), 1298. doi: 10.3390/s16081298

Gómez-Rodríguez, C., Díaz-Paniagua, C., Serrano, L., Florencio, M., & Portheault, A. (2009). Mediterranean temporary ponds as amphibian breeding habitats: the importance of preserving pond networks. Aquatic Ecology, 43, 1179–1191. doi: 10.1007/s10452-009-9235-x

Gxokwe, S., Dube, T., Mazvimavi, D., & Grenfell, M. (2022). Using cloud computing techniques to monitor long-term variations in ecohydrological dynamics of small seasonally-flooded wetlands in semi-arid South Africa. Journal of Hydrology, 612, 128080. doi: 10.1016/j.jhydrol.2022.128080

Haralick, R. M., Shanmugam, K., & Dinstein, I. H. (1973). Textural features for image classification. IEEE Transactions on systems, man, and cybernetics, (6), 610-621. doi: 10.1109/TSMC.1973.4309314

Halabisky, M., Moskal, L. M., Gillespie, A., & Hannam, M. (2016). Reconstructing semi-arid wetland surface water dynamics through spectral mixture analysis of a time series of Landsat satellite images (1984– 2011). Remote sensing of environmen,t 177, 171–183. doi: 10.1016/j.rse.2016.02.040

Herceg-Szórádi, Z., Demeter, L., & Csergő, A. M. (2023). Small area and low connectivity constrain the diversity of plant life strategies in temporary ponds. Diversity and Distributions. doi: 10.1111/ddi.13685

Huang, W., DeVries, B., Huang, C., Lang, M. W., Jones, J. W., Creed, I. F., & Carroll, M. L. (2018). Automated extraction of surface water extent from Sentinel-1 data. Remote Sensing, 10(5), 797. doi: 10.3390/rs10050797

Hoekstra, A. Y. (2009). Human appropriation of natural capital: A comparison of ecological footprint and water footprint analysis. Ecological economics, 68(7), 1963–1974. doi: 10.1016/j.ecolecon.2008.06.021

Institute for European Environmental Policy, Poláková, J., Tucker, G., Hart, K., Dwyer, J., & Rayment, M. (2011). Addressing biodiversity and habitat preservation through measures applied under the Common Agricultural Policy. London: Institute for European Environmental Policy.

Liaw, A., & Wiener, M. (2002). Classification and regression by randomForest. R news, 2(3), 18–22.

Liu, Z., Blasch, E., Bhatnagar, G., John, V., Wu, W., & Blum, R. S. (2018). Fusing synergistic information from multi-sensor images: an overview from implementation to performance assessment. Information Fusion, 42, 127–145. doi: 10.1016/j.inffus.2017.10.010

Jeffrey, P., & Gearey, M. (2006). Integrated water resources management: lost on the road from ambition to realisation?. Water science and technology, 53(1), 1–8. doi: 10.2166/wst.2006.001

Jenkins, K., & Warren, R. (2015). Quantifying the impact of climate change on drought regimes using the Standardised Precipitation Index. Theoretical and Applied Climatology, 120, 41–54. doi: 10.1007/s00704-014-1143-x

Jiang, Z., Wen, Y., Zhang, G., & Wu, X. (2022). Water Information Extraction Based on Multi-Model RF Algorithm and Sentinel-2 Image Data. Sustainability, 14(7), 3797. doi: 10.3390/su14073797

Jiang, W., Ni, Y., Pang, Z., He, G., Fu, J., Lu, J., et al (2020). A new index for identifying water body from sentinel-2 satellite remote sensing imagery. ISPRS Annals of Photogrammetry, Remote Sensing & Spatial Information Sciences, (3).

Klein, I., Gessner, U., Dietz, A. J., & Kuenzer, C. (2017). Global WaterPack–A 250 m resolution dataset revealing the daily dynamics of global inland water bodies. Remote sensing of environment, 198, 345-362. doi: 10.1016/j.rse.2017.06.045

Mahdianpari, M., Salehi, B., Mohammadimanesh, F., Homayouni, S., & Gill, E. (2018). The first wetland inventory map of newfoundland at a spatial resolution of 10 m using sentinel-1 and sentinel-2 data on the google earth engine cloud computing platform. Remote Sensing, 11(1), 43. doi: https://www.mdpi.com/2072-4292/11/1/43 10.3390/rs11010043

Mayer, T., Poortinga, A., Bhandari, B., Nicolau, A. P., Markert, K., Thwal, N. S., et al (2021). Deep learning approach for Sentinel-1 surface water mapping leveraging Google Earth Engine. ISPRS Open Journal of Photogrammetry and Remote Sensing, 2, 100005. doi: 10.1016/j.ophoto.2021.100005

Mapper, L., & Plus, E. T. M. Landsat Collection 2 Level-3 Dynamic Surface Water Extent Science Product.

Martensen, A. C., Saura, S., & Fortin, M. J. (2017). Spatio-temporal connectivity: assessing the amount of reachable habitat in dynamic landscapes. Methods in Ecology and Evolutio,n 8(10), 1253–1264. doi: 10.1111/2041-210X.12799

Mondejar, J. P., & Tongco, A. F. (2019). Near infrared band of Landsat 8 as water index: a case study around Cordova and Lapu-Lapu City, Cebu, Philippines. Sustainable Environment Research, 29, 1–15. doi: 10.1186/s42834-019-0016-5

Morante-Carballo, F., Montalván-Burbano, N., Quiñonez-Barzola, X., Jaya-Montalvo, M., & Carrión-Mero, P. (2022). What do we know about water scarcity in semi-arid zones? A global analysis and research trends. Water, 14(17), 2685. doi: 10.3390/w14172685

McKenzie, L. J., Finkbeiner, M. A., & Kirkman, H. (2001). Methods for mapping seagrass distribution. Global seagrass research methods, 101–121. doi: 10.1016/B978-044450891-1/50006-2

McNairn, H., & Brisco, B. (2004). The application of C-band polarimetric SAR for agriculture: A review. Canadian Journal of Remote Sensing, 30(3), 525–542. doi: 10.5589/m03-069

Millard, K., & Richardson, M. (2015). On the importance of training data sample selection in random forest image classification: A case study in peatland ecosystem mapping. Remote sensing, 7(7), 8489–8515. doi: 10.3390/rs70708489

Nasirzadehdizaji, R., Balik Sanli, F., Abdikan, S., Cakir, Z., Sekertekin, A., & Ustuner, M. (2019). Sensitivity analysis of multi-temporal Sentinel-1 SAR parameters to crop height and canopy coverage. Applied Sciences, 9(4), 655. doi: 10.3390/app9040655

Olmo, C., Galvez, A., Bisquert-Ribes, M., Bonilla, F., Vega, C., Castillo-Escriva, A., et al (2022). The environmental framework of temporary ponds: A tropical-mediterranean comparison. Catena, 210, 105845. doi: 10.1016/j.catena.2021.105845

Ozesmi, S. L., & Bauer, M. E. (2002). Satellite remote sensing of wetlands. Wetlands ecology and management, 10, 381–402. doi: 10.1023/A:1020908432489

Peña-Luque, S., Ferrant, S., Cordeiro, M. C., Ledauphin, T., Maxant, J., & Martinez, J. M. (2021). Sentinel-1&2 multitemporal water surface detection accuracies, evaluated at regional and reservoirs level. Remote Sensing, 13(16), 3279. doi: 10.3390/rs13163279

Pekel, J. F., Cottam, A., Gorelick, N., & Belward, A. S. (2016). High-resolution mapping of global surface water and its long-term changes. Nature, 540(7633), 418–422. doi: 10.1038/nature20584

Petropoulos, G. P., Ireland, G., & Barrett, B. (2015). Surface soil moisture retrievals from remote sensing: Current status, products & future trends. *Physics and Chemistry of the Earth*, Parts A/B/C, 83, 36–56. doi: 10.1016/j.pce.2015.02.009

Prošek, J., Gdulová, K., Barták, V., Vojar, J., Solský, M., Rocchini, D., & Moudrý, V. (2020). Integration of hyperspectral and LiDAR data for mapping small water bodies. International Journal of Applied Earth Observation and Geoinformation, 92, 102181. doi: 10.1016/j.jag.2020.102181

Rachid, N., Issam, N., Abdelkrim, B., Abdelhamid, A., & Amina, H. (2022). Pond Energy Dynamics, Evaporation Rate and Ensemble Deep Learning Evaporation Prediction: Case Study of the Thomas Pond— Brenne Natural Regional Park (France). Water, 14(6), 923. doi: 10.3390/w14060923

Radočaj, D., Obhođaš, J., Jurišić, M., & Gašparović, M. (2020). Global open data remote sensing satellite missions for land monitoring and conservation: A review. Land, 9(11), 402. doi: 10.3390/land9110402

Radoux, J., Chomé, G., Jacques, D. C., Waldner, F., Bellemans, N., Matton, N., et al (2016). Sentinel-2’s potential for sub-pixel landscape feature detection. Remote Sensing, 8(6), 488. doi: 10.3390/rs8060488

Reid, A. J., Carlson, A. K., Creed, I. F., Eliason, E. J., Gell, P. A., Johnson, P. T., et al (2019). Emerging threats and persistent conservation challenges for freshwater biodiversity. Biological Reviews, 94(3), 849–873. doi: 10.1111/brv.12480

Richardson, A. J., & Wiegand, C. L. (1977). Distinguishing vegetation from soil background information. Photogrammetric engineering and remote sensing, 43(12), 1541–1552.

Ritchie, J. C., Zimba, P. V., & Everitt, J. H. (2003). Remote sensing techniques to assess water quality. Photogrammetric engineering & remote sensing, 69(6), 695–704. doi: 10.14358/PERS.69.6.695

Rouse, J. W., Haas, R. H., Schell, J. A., & Deering, D. W. (1974). Monitoring vegetation systems in the Great Plains with ERTS. NASA Spec. Publ, 351(1), 309.

Samaniego, L., Thober, S., Kumar, R., Wanders, N., Rakovec, O., Pan, M., et al. (2018). Anthropogenic warming exacerbates European soil moisture droughts. Nature Climate Change, 8(5), 421–426. doi: 10.1038/s41558-018-0138-5

Sawaya, K. E., Olmanson, L. G., Heinert, N. J., Brezonik, P. L., & Bauer, M. E. (2003). Extending satellite remote sensing to local scales: land and water resource monitoring using high-resolution imagery. Remote sensing of Environment, 88(1-2), 144–156. doi: 10.1016/j.rse.2003.04.006

Sethre, P. R., Rundquist, B. C., & Todhunter, P. E. (2005). Remote detection of prairie pothole ponds in the Devils Lake Basin, North Dakota. GIScience & Remote Sensing, 42(4), 277–296. doi: 10.2747/1548-1603.42.4.277

Silva, J. P., Gameiro, J., Valerio, F., & Marques, A. T. (2024). Portugal’s farmland bird crisis requires action. Science, 383(6679), 157–157. doi: 10.1126/science.adn1390

Slagter, B., Tsendbazar, N. E., Vollrath, A., & Reiche, J. (2020). Mapping wetland characteristics using temporally dense Sentinel-1 and Sentinel-2 data: A case study in the St. Lucia wetlands, South Africa. International Journal of Applied Earth Observation and Geoinformatio, 8n6, 102009. doi: 10.1016/j.jag.2019.102009

Sebastián-González, E., & Green, A. J. (2014). Habitat use by waterbirds in relation to pond size, water depth, and isolation: lessons from a restoration in southern Spain. Restoration Ecology, 22(3), 311–318. doi: 10.1111/rec.12078

Serrano, L., Ustin, S. L., Roberts, D. A., Gamon, J. A., & Penuelas, J. (2000). Deriving water content of chaparral vegetation from AVIRIS data. Remote sensing of Environment, 74(3), 570–581. doi: 10.1016/S0034-4257(00)00147-4

Scholes, R. J. (2020). The future of semi-arid regions: A weak fabric unravels. Climate, 8(3), 43. doi: 10.3390/cli8030043

Semeniuk, C. A., & Semeniuk, V. (1995). A geomorphic approach to global classification for inland wetlands. Classification and Inventory of the World’s Wetlands, 103-124.

Shen, X., Wang, D., Mao, K., Anagnostou, E., & Hong, Y. (2019). Inundation extent mapping by synthetic aperture radar: A review. Remote Sensing, 11(7), 879. doi: 10.3390/rs11070879

Silva, J. P., Catry, I., Palmeirim, J. M., & Moreira, F. (2015). Freezing heat: thermally imposed constraints on the daily activity patterns of a free-ranging grassland bird. Ecosphere, 6(7), 1–13. doi: 10.1890/ES14-00454.1

Soria, J., Ruiz, M., & Morales, S. (2022). Monitoring Subaquatic Vegetation Using Sentinel-2 Imagery in Gallocanta Lake (Aragón, Spain). Earth, 3(1), 363–382. doi: 10.3390/earth3010022

Stehman, S. V. (1997). Selecting and interpreting measures of thematic classification accuracy. Remote sensing of Environment, 62(1), 77–89. doi: 10.1016/S0034-4257(97)00083-7

Sun, F., Sun, W., Chen, J., & Gong, P. (2012). Comparison and improvement of methods for identifying waterbodies in remotely sensed imagery. International journal of remote sensin,g 33(21), 6854–6875. doi: 10.1080/01431161.2012.692829

Swets, J. A. (1988). Measuring the accuracy of diagnostic systems. Science, 240(4857), 1285–1293. doi: 10.1126/science.3287615

Tang, H., Lu, S., Ali Baig, M. H., Li, M., Fang, C., & Wang, Y. (2022). Large-scale surface water mapping based on landsat and sentinel-1 images. Water, 14(9), 1454. doi: 10.3390/w14091454

Team, R. C. (2021). R: A language and environment for statistical computing. Published online 2020.

Torres, R., Snoeij, P., Geudtner, D., Bibby, D., Davidson, M., Attema, E., et al (2012). GMES Sentinel-1 mission. Remote sensing of environment, 120, 9–24. doi: 10.1016/j.rse.2011.05.028

Tucker, M. A., Böhning-Gaese, K., Fagan, W. F., Fryxell, J. M., Van Moorter, B., Alberts, S. C., et al (2018). Moving in the Anthropocene: Global reductions in terrestrial mammalian movements. Science, 359(6374), 466–469. doi: 10.1126/science.aam9712

Tulbure, M. G., Broich, M., Stehman, S. V., & Kommareddy, A. (2016). Surface water extent dynamics from three decades of seasonally continuous Landsat time series at subcontinental scale in a semi-arid region. Remote Sensing of Environment, 178, 142–157. doi: 10.1016/j.rse.2016.02.034

Tulbure, M. G., Broich, M., Perin, V., Gaines, M., Ju, J., Stehman, S. V., et al (2022). Can we detect more ephemeral floods with higher density harmonized Landsat Sentinel 2 data compared to Landsat 8 alone?. ISPRS Journal of Photogrammetry and Remote Sensing, 185, 232–246. doi: 10.1016/j.isprsjprs.2022.01.021

United Na ons. Department of Economic and Social Affairs. (2022). *The Sustainable Development Goals: Report* 2022. UN.

Vanderhoof, M. K., Alexander, L., Christensen, J., Solvik, K., Nieuwlandt, P., & Sagehorn, M. (2023). High-frequency time series comparison of Sentinel-1 and Sentinel-2 satellites for mapping open and vegetated water across the United States (2017–2021). Remote Sensing of Environment, 288, 113498. doi: 10.1016/j.rse.2023.113498

Valavi, R., Elith, J., Lahoz-Monfort, J. J., & Guillera-Arroita, G. (2018). blockCV: An r package for generating spatially or environmentally separated folds for k-fold cross-validation of species distribution models. Biorxiv, 357798.

Valerio, F., Ferreira, E., Godinho, S., Pita, R., Mira, A., Fernandes, N., & Santos, S. M. (2020). Predicting microhabitat suitability for an endangered small mammal using sentinel-2 data. Remote Sensing, 12(3), 562. doi: https://www.mdpi.com/2072-4292/12/3/562 10.3390/rs12030562

Valerio, F., Godinho, S., Marques, A. T., Crispim-Mendes, T., Pita, R., & Silva, J. P. (2024). GEE_xtract: High-quality remote sensing data preparation and extraction for multiple spatio-temporal ecological scaling. Ecological Informatics, 102502(80), 1574–9541. doi: 10.1016/j.ecoinf.2024.102502

Vasudev, D., Fletcher Jr, R. J., Goswami, V. R., & Krishnadas, M. (2015). From dispersal constraints to landscape connectivity: lessons from species distribution modeling. Ecography, 38(10), 967–978. doi: 10.1111/ecog.01306

Veldkamp, T. I., Wada, Y., de Moel, H., Kummu, M., Eisner, S., Aerts, J. C., & Ward, P. J. (2015). Changing mechanism of global water scarcity events: Impacts of socioeconomic changes and inter-annual hydro-climatic variability. Global Environmental Change, 32, 18–29. doi: 10.1016/j.gloenvcha.2015.02.011

Vreugdenhil, M., Navacchi, C., Bauer-Marschallinger, B., Hahn, S., Steele-Dunne, S., Pfeil, I., et al (2020). Sentinel-1 cross ratio and vegetation optical depth: A comparison over Europe. Remote Sensing, 12(20), 3404. doi: 10.3390/rs12203404

Wang, L., Bie, W., Li, H., Liao, T., Ding, X., Wu, G., & Fei, T. (2022). Small Water Body Detection and Water Quality Variations with Changing Human Activity Intensity in Wuhan. Remote Sensing, 14(1), 200. doi: 10.3390/rs14010200

Warton, D. I., & Weber, N. C. (2002). Common slope tests for bivariate errors-in-variables models. Biometrical Journal: Journal of Mathematical Methods in Biosciences, 44(2), 161–174. doi: 10.1002/1521-4036(200203)44:2%3C161::AID-BIMJ161%3E3.0.CO;2-N

Wickens, G.E. (1998). Arid and Semi-arid Environments of the World. In: Ecophysiology of Economic Plants in Arid and Semi-Arid Lands. Adaptations of Desert Organisms. Springer, Berlin, Heidelberg. doi: 10.1007/978-3-662-03700-3_2

Williams, P., Whitfield, M., Biggs, J., Bray, S., Fox, G., Nicolet, P., & Sear, D. (2004). Comparative biodiversity of rivers, streams, ditches and ponds in an agricultural landscape in Southern England. Biological conservation, 115(2), 329–341. doi:∼ 10.1016/S0006-3207(03)00153-8

Wilson, E. H., & Sader, S. A. (2002). Detection of forest harvest type using multiple dates of Landsat TM imagery. Remote Sensing of Environment, 80(3), 385–396. doi: 10.1016/S0034-4257(01)00318-2

Wulder, M. A., & Coops, N. C. (2014). Satellites: Make Earth observations open access. Nature, 513(7516), 30–31. doi: 10.1038/513030a

Zamora-Marín, J. M., Zamora-López, A., Oliva-Paterna, F. J., Torralva, M., Sánchez-Montoya, M. M., & Calvo, J. F. (2024). From small waterbodies to large multi-service providers: Assessing their ecological multifunctionality for terrestrial birds in Mediterranean agroecosystems. *Agriculture*, Ecosystems & Environment, 359, 108760. doi: 10.1016/j.agee.2023.108760

Zeller, K. A., Lewison, R., Fletcher Jr, R. J., Tulbure, M. G., & Jennings, M. K. (2020). Understanding the importance of dynamic landscape connectivity. Land, 9(9), 303. doi: 10.3390/land9090303

Zhou, Y., Dong, J., Xiao, X., Xiao, T., Yang, Z., Zhao, G., et al (2017). Open surface water mapping algorithms: A comparison of water-related spectral indices and sensors. Water, 9(4), 256. doi: 10.3390/w9040256

Zou, Z., Xiao, X., Dong, J., Qin, Y., Doughty, R. B., Menarguez, M. A., et al (2018). Divergent trends of open-surface water body area in the contiguous United States from 1984 to 2016. Proceedings of the National Academy of Sciences, 115(15), 3810–3815. doi: 10.1073/pnas.1719275115

