## Supplementary material for "Multi-Temporal Remote Sensing of Inland Surface Waters: A Fusion of Sentinel-1&2 Data Applied to Small Seasonal Ponds in Semiarid Environments": SM

Supplementary Tables

The dataset referring to maximum water storage capacity was developed using imageries present in the Google Earth software (Google), and finalized through the QGIS software (QGIS Development Team, 2009), using a reference period between 2017 and 2021, as detailed below.

| **Study Areas** | **Identified water bodies** | **Satellite timestamp** |
| --- | --- | --- |
| **PT2** | 816 | 18/11/2017 |
| **PT1** | 185 | 19/11/2017 |
| **SP1** | 1135 | 19/08/2019 |
| **SP1** | 1220 | 20/08/2019 |
| **PT2** | 8 | 31/03/2021 |
| **PT2** | 2 | 17/09/2021 |

*Table SM1* Dataset referring to the characterization of water bodies with maximum water storage capacity.

The dataset referring to minimum water storage capacity was developed using a reference period in 2021, as detailed below.

| **Study Areas** | **Identified water bodies** | **Satellite timestamp** |
| --- | --- | --- |
| **PT2** | 2 | 31/03/2021 |
| **PT1** | 122 | 15/05/2021 |
| **PT2** | 480 | 15/05/2021 |
| **PT2** | 2 | 17/09/2021 |

*Table SM2* Dataset referring to the characterization of water bodies with effective water surface.

| Metric | Type | Unit | Range | SD | SE | Score |
| --- | --- | --- | --- | --- | --- | --- |
| OOB error rate | Error | Percentage | 0 to 100 | 0.646 | 0.723 | 26.63% |
| Sensitivity | Accuracy |  |  | 0.04 | 0.03 | 69.40% |
| Specificity | Accuracy |  |  | 0.04 | 0.04 | 75.11% |
| Accuracy | Accuracy |  |  | 0.04 | 0.04 | 72.56% |
| AUC | Accuracy |  |  | 0.04 | 0.04 | 72.70% |
| F1 | Accuracy |  |  | 0.05 | 0.02 | 69.85% |
| MCC | Accuracy | Correlation coefficient | -1 to 1 | 0.06 | 0.02 | 0.44 |

*Table SM3* Metrics scores (OOB error rate, Sensitivity, Specificity, Accuracy, AUC, F1, MCC) defining RF model performances for surface water occurrence classification, without considering the Sentinel-1 data. Scores are coupled with standard deviation (SD) and standard error (SD), and averaged on the basis of the geographically distinct 5-fold RF cross-validations.

Supplementary Figures

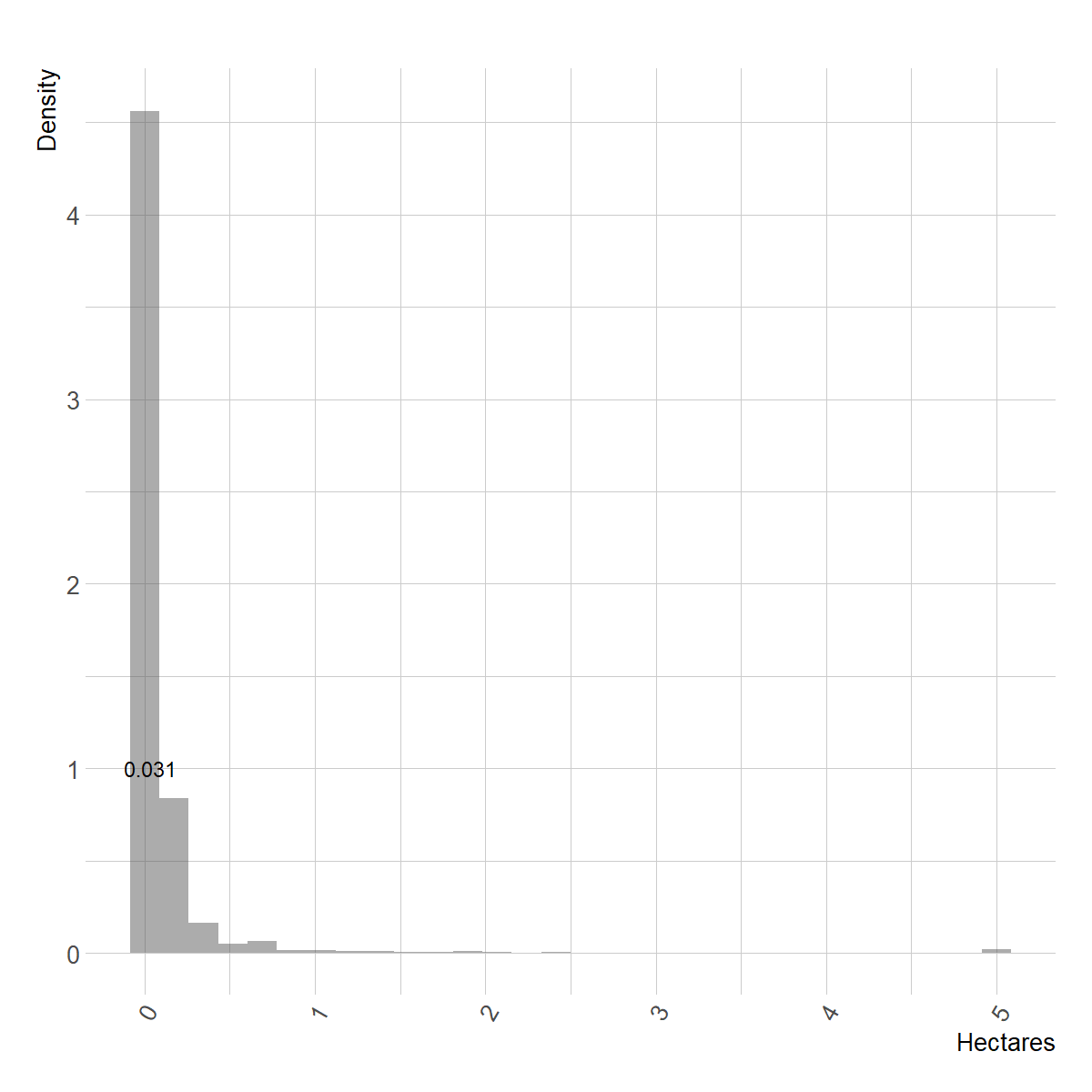

*Figure SM1* Density plot of hectares values and mean distribution, from the 3366 sampled water bodies within the three study areas. The interval values are fixed between 0 and 5 ha according to the definition of small water bodies (ponds) defined by De Meester et al., (2005). The score in the density plot represents the median value of the distribution.

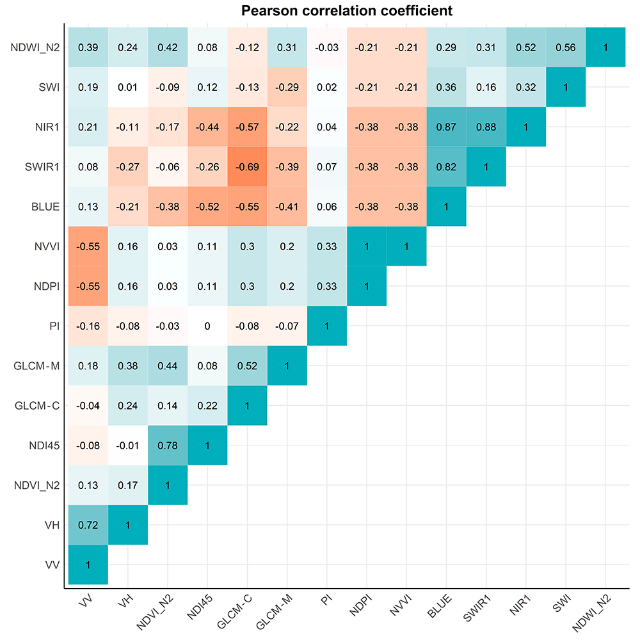

*Figure SM2* Upper-triangular correlation matrix (Pearson correlation coefficients) including the 14 retained Sentinel1&2 derived features.

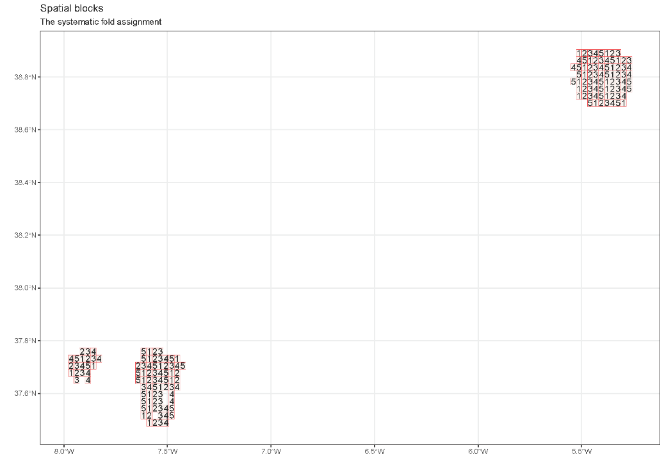

*Figure SM3* The spatial blocking strategy is outlined in red on the basis of the 4x4km grid size (Figure 1), where in each block is numerically allocated a fold used for spatial cross-validation analysis.

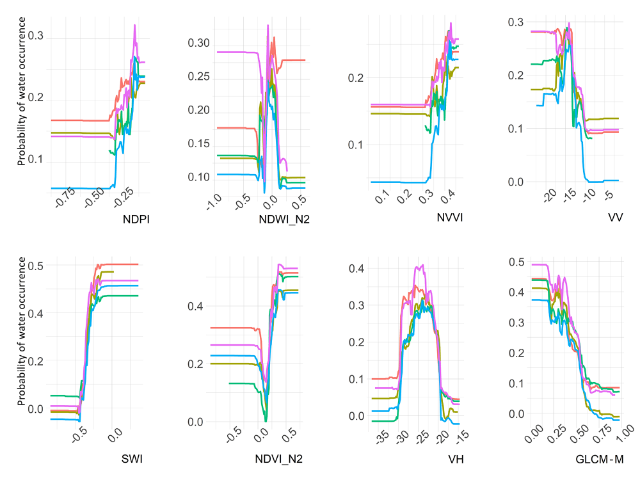

*Figure SM4* Curve responses of remaining features explaining water occurrence. For each feature, several curves are depicted, in turn reflecting a fold of the 5 RF cross-validation analyses.

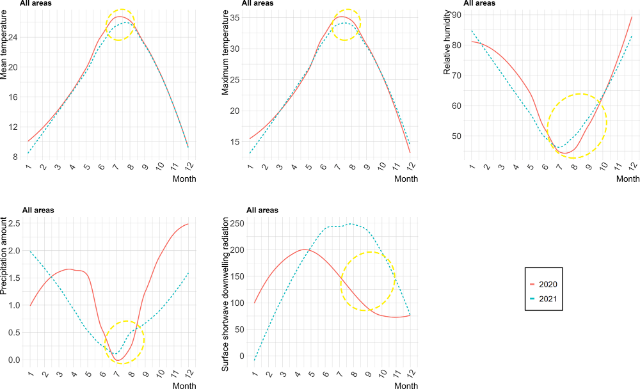

*Figure SM5* Smoothed data of the E-OBS time series for mean temperature, maximum temperature, precipitation amount, relative humidity, and surface shortwave downwelling radiation. The continuous red color line represents the 2020 trend, whilst the 2021 is represented by the blue and short-dashed line. The yellow long-dashed circle highlights the difference in trends between years, approximately focusing on the period interval when anomalies in water surface extent were evidenced (September-October; Figure 7).

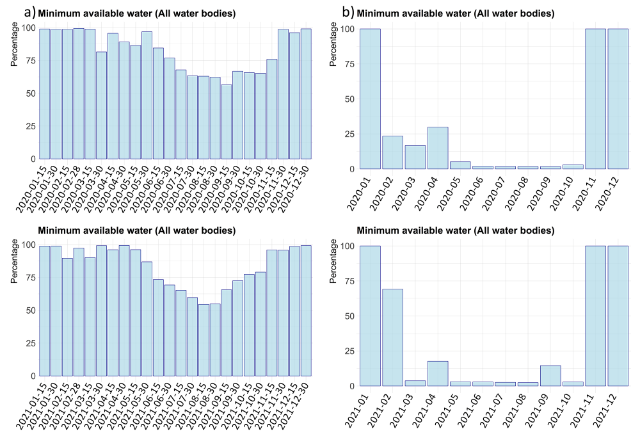

*Figure SM6* Dynamic surface water occurrence expressed as proportion (percentage) of water bodies with minimum available water, considering the three semiarid regions collectively. For each date, a bar illustrates the percentage of water bodies with water. The time series are produced by SLSW (a) and LGSW (b) models with an interval of 15 days and 30 days, respectively. The forecasting time series were reproduced for the year 2020 (upper panels) and 2021 (lower panels).

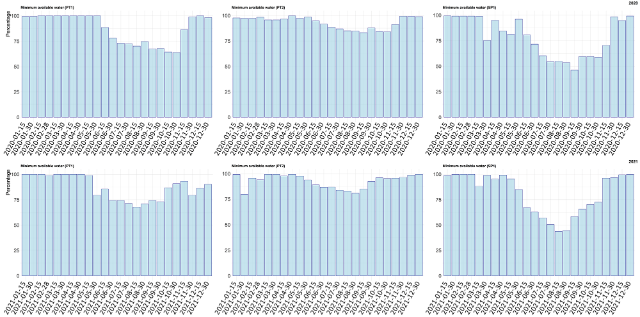

*Figure SM7* Dynamic surface water occurrence expressed as proportion (percentage) of water bodies with minimum available water, considering the three semiarid regions (PT1, PT2, and SP1) separately. For each date, a bar illustrates the percentage of water bodies with water. The time series are produced by the SLSW product with an interval of 15 days. The time series was reproduced for the year 2020 (upper panels) and 2021 (lower panels).

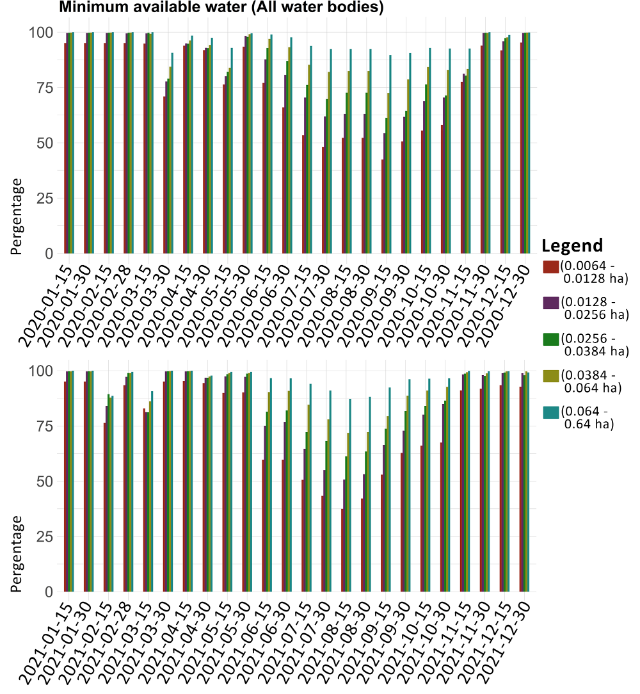

*Figure SM8* Dynamic surface water occurrence expressed as proportion (percentage), with boxplots representing different water body sizes (0.0064-0.0128, 2 pixels; 0.0128-0.0256, 3 pixels; 0.0256-0.0384, 5 pixels; 0.0384-0.064, 10 pixels; 0.064-0.64, 100 pixels). The three semiarid regions were considered collectively and for each date, a bar illustrates the percentage of water bodies with water. The time series represent the SLSW product, and were reproduced for the years 2020 (upper panels) and 2021 (lower panels).

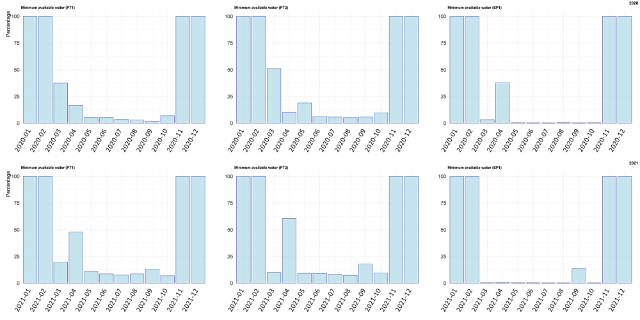

*Figure SM9* Dynamic surface water occurrence expressed as proportion (percentage) of water bodies with minimum available water, considering the three semiarid regions (PT1, PT2, and SP1) separately. For each date, a bar illustrates the percentage of water bodies with water. The time series are produced by the LGSW product with an interval of 15 days. The time series was reproduced for the year 2020 (upper panels) and 2021 (lower panels).

Supplementary Material 1.1: Optimal threshold calculation

The optimal threshold to discriminate water vs non-water classes was based on 5 approaches aggregating predicted values, as well contrasting predicted values with those measured, or by taking a fixed value. The first criteria consisted in minimizing the difference between the sensitivity (true positive rate) and specificity (true negative rate), namely where positive observations are just as likely to be wrong as negative observations (Fielding & Bell, 1997). The second criteria consisted in maximizing the sum of sensitivity and specificity (Cantor et al., 1999), namely by minimizing the mean of the error rate for positive observations and the error rate for negative observations. The third criteria consisted in using the original species prevalence as threshold (Cramer, 2003). The fourth criteria consisted in taking the mean of the probabilities of occurrence of occupied locations for presence/absence data as the threshold (Cramer, 2003). The last criteria utilized a fixed threshold of 0.5 (Manel et al., 1999; Nenzén & Araújo, 2011). The calculation of optimal threshold was conducted in R statistical environment (v.4.2.0; R Core Team, 2021) utilizing the ‘*PresenceAbsence*’ (v.1.1.10) package (Freeman & Moisen, 2008).

REFERENCES:

Cantor, S. B., Sun, C. C., Tortolero-Luna, G., Richards-Kortum, R., & Follen, M. (1999). A comparison of C/B ratios from studies using receiver operating characteristic curve analysis. *Journal of clinical epidemiology*, *52*(9), 885-892.

Cramer, J. S. (2003). *Logit models from economics and other fields*. Cambridge University Press.

De Meester, L., Declerck, S., Stoks, R., Louette, G., Van De Meutter, F., De Bie, T., ... & Brendonck, L. (2005). Ponds and pools as model systems in conservation biology, ecology and evolutionary biology. *Aquatic conservation: Marine and freshwater ecosystems*, *15*(6), 715-725.

Google, Inc., Google Earth software, http://earth.google.com/ [last accessed on March 13, 2022]

Fielding, A. H., & Bell, J. F. (1997). A review of methods for the assessment of prediction errors in conservation presence/absence models. *Environmental conservation*, *24*(1), 38-49.

Freeman, E. A., & Moisen, G. (2008). PresenceAbsence: An R package for presence absence analysis. Journal of Statistical Software. 23 (11): 31 p. Manel, S., Dias, J. M., & Ormerod, S. J. (1999). Comparing discriminant analysis, neural networks and logistic regression for predicting species distributions: a case study with a Himalayan river bird. *Ecological modelling*, *120*(2-3), 337-347.

Nenzén, H. K., & Araújo, M. B. (2011). Choice of threshold alters projections of species range shifts under climate change. *Ecological Modelling*, *222*(18), 3346-3354.

QGIS Development Team, 2009. QGIS Geographic Information System. Open Source Geospatial Foundation. URL http://qgis.org

Team, R. C. (2021). R: A language and environment for statistical computing. Published online 2020.
